# A mass spectrometry-based method for the determination of *in vivo* biodistribution of tumor targeting small molecule-metal conjugates

**DOI:** 10.1101/2022.03.04.483029

**Authors:** Ettore Gilardoni, Aureliano Zana, Andrea Galbiati, Theo Sturm, Jacopo Millul, Samuele Cazzamalli, Dario Neri, Riccardo Stucchi

## Abstract

Nuclear medicine plays a key role in modern diagnosis and cancer therapy. The development of tumor targeting radionuclide conjugates (also named Small Molecule-Radio Conjugates - SMRCs) represents a significant improvement over the clinical use of metabolic radiotracers (e.g., [^18^F]-Fluorodeoxyglucose) for imaging and over the application of biocidal external beam radiations for therapy. During the discovery of SMRCs, molecular candidates must be carefully evaluated typically by performing biodistribution assays in preclinical tumor models. Quantification methodologies based on radioactive counts are typically demanding due to safety concerns, availability of radioactive material, and infrastructures. In this article, we report the development of a mass spectrometry (MS)-based method for the detection and quantification of small molecule-metal conjugates (SMMCs) as cold surrogates of SMRCs. We applied this methodology for the evaluation of the biodistribution of a particular class of tumor-targeting drug candidates based on ^nat^Lu, ^nat^Ga, ^nat^F and directed against Fibroblast Activation Protein (FAP). The reliability of the LC-MS analysis was validated by direct comparison of MS-based and radioactivity-based biodistribution data. Results show that MS biodistribution of stable isotope metal conjugates is an orthogonal tool for the preclinical characterization of different classes of radiopharmaceuticals.

## Introduction

Small molecule-radio conjugates (SMRCs) are radioactive compounds which typically consist of two components: a radionuclide included in a molecular structure and a targeting moiety which is responsible for specific accumulation of the radioactive payload at the site of disease (typically cancer lesions)^1^. Thanks to this peculiar design, SMRCs find applications both as diagnostic tracers and as radiotherapeutic agents in modern nuclear medicine. In the recent years, clinical success of Lutathera®^2^ and of PSMA-617^3^ for the treatment of patients with neuroendocrine tumors and metastatic castration resistant prostate cancer has demonstrated the potential of this class of theragnostic. Many novel SMRC drug candidates are being developed and studied in clinical trials for diagnostic and therapeutic applications of patients with cancer and other types of chronic malignancies^4,5^,

Among several new tumor-targeting radiopharmaceuticals, SMRCs targeting fibroblast activation protein (FAP) are promising pan-tumoral compounds which accumulate to tumors with exquisite selectivity against normal tissues^4,6,7^. FAP is a membrane-bound serine protease abundantly expressed on the stroma of most epithelial cancers^8^. FAP has been validated as high-quality tumor-associated antigen by immunohistochemistry and nuclear medicine ^4,6,7,9^. Our group has recently developed OncoFAP, a small molecule targeting FAP with ultra-high affinity (K_D_= 680 pM), able to selectively deliver radionuclides, such as Lutetium and Gallium, to the tumor site^10^. A Gallium-68 OncoFAP derivative shows excellent tumor-targeting properties in patients with solid malignancies, with tumor-to-organ ratio of ~20-to-1 at early time points^6^. This compound represents an ideal candidate for imaging applications. More recently, we developed a dimeric derivative owing two targeting moieties (i.e., BiOncoFAP), showing higher tumor uptake and residence time than OncoFAP in preclinical models of cancer^11^.

The biodistribution properties of novel tumor targeting SMRCs are of key importance for the success of the diagnosis and therapeutic outcome. Preclinical biodistribution of radiopharmaceuticals in tumor-bearing animals is typically evaluated by direct measurement of the radioactivity present in each organ after sacrifice of experimental animals^10,11^. To date, these radioactivity-based analyses are still crucial to evaluate the tumor accumulation and to obtain spatial biodistribution data of the compounds in the animal/human body. On the contrary, for most of the other non-radioactive pharmaceuticals, LC-MS analysis represents the analytical method of choice when it comes to accurately quantify the analyte of interest in biological specimens^12,13^. Since radioactivity-based experiments are challenging because of safety reasons (i.e., potential exposure of the operator to harmful radiation doses) and availability of dedicated infrastructures, the development of an alternative analytical methodology would open exciting new possibilities.

To our knowledge, most of the studies reported in the literature for the analysis of nonradioactive metal-conjugates rely on HPLC-UV or ICP-MS methods^14^, while only a very limited number exploit LC-MS techniques^15,16^.

In this work, we developed a MS-based quantification methodology which allows to determine quantitative biodistribution of Small molecule-metal conjugates (SMMCs) surrogates of SMRCs based on cold non-radioactive isotopes, aiming at offering a valid alternative to the use of hot radionuclides throughout the discovery and development of radiopharmaceuticals. By comparing “cold” biodistribution data (i.e., data obtained after administration of SMMCs) obtained by mass spectrometry and radio-biodistribution data (i.e., data obtained after administration of the corresponding SMRCs), we validated this novel LC-MS method as an orthogonal, safe, green, and easy-to-implement option to classical radioactivity-based methodologies.

## Experimental Section

### Chemicals and Materials

All reagents and solvents were purchased from Sigma Aldrich, VWR, Combi-Blocks, CheMatech and used as supplied.

### Chemical Synthesis

OncoFAP and BiOncoFAP, as well as their metal chelator conjugates and “cold”-labeled derivatives were synthesized through established protocols^6,10,11^.

For the synthesis of internal standards (ISs), we used isotopically labeled building blocks. In particular, ^13^C4-succinic anhydride was exploited for OncoFAP derivatives, while ^13^C6^15^N_2-L_-Lysine was used for BiOncoFAP-conjugates.

Detailed experimental chemical procedures are described in the Supplemental Informations.

### Animal studies

All animal experiments were conducted in accordance with Swiss animal welfare laws and regulations under the license number ZH006/2021 granted by the Veterinäramt des Kantons Zürich.

### In vivo biodistribution of OncoFAP molecules in tumor-bearing mice

Tumor cells (HT-1080.hFAP, or SK-RC-52.hFAP cells) were grown to 80% confluence and detached with Trypsin-EDTA 0.05% (Gibco). Cells were then resuspended in Hanks’ Balanced Salt Solution medium (Gibco). Aliquots of 5 to 10 × 10^6^ cells (100 to 150 μL of suspension) were injected subcutaneously in the right flanks of female athymic Balb/c AnNRj-Foxn1 mice (6 to 8 weeks of age, Janvier).

Mice bearing subcutaneous HT-1080.hFAP tumors were injected intravenously with [^nat^Lu]Lu-OncoFAP-DOTAGA, [^nat^Lu]Lu-BiOncoFAP-DOTAGA, [^nat^Ga]Ga-OncoFAP-DOTAGA, or [^nat^Ga]Ga-BiOncoFAP-DOTAGA (5 nmol dissolved in sterile PBS, pH 7.4). Animals were sacrificed 1 h after intravenous injection. Fresh blood was collected in lithium heparin tubes (BD Microcontainer LH Tubes), vortexed and centrifuged (15’000 g, 15 min). Plasma was frozen and stored at −80 °C. Healthy organs and tumors were subsequently excised, frozen with dry ice and stored at −80 °C.

Mice bearing subcutaneous SK-RC-52.hFAP tumors were injected intravenously with [^nat^F]AlF-OncoFAP-NODAGA, or [^nat^F]AlF-OncoFAP-NOTA, (10 nmol dissolved in sterile PBS, pH 7.4). Animals were sacrificed 2 h after intravenous injection. Fresh blood was collected in lithium heparin tubes (BD Microcontainer LH Tubes), vortexed and centrifuged (15’000 g, 15 min). Plasma was frozen and stored at −80 °C. Healthy organs and tumors were subsequently excised, frozen with dry ice and stored at −80 °C

### Sample preparation for Mass Spectrometry analysis

50 mg of mice tissues were resuspended in 600 μL of a solution containing 95 % ACN and 0.1% FA to induce protein precipitation. In parallel, 50 μL of a solution 600 nM of internal standard ([^nat^Lu]Lu-^13^C4-OncoFAP-DOTAGA, or [^nat^Lu]Lu-^13^C6^15^N_2_-BiOncoFAP-DOTAGA, or [^nat^Ga]Ga-^13^C4-OncoFAP-DOTAGA, or [^nat^Ga]Ga-^13^C6^15^N_2_-BiOncoFAP-DOTAGA, or [^nat^F]AlF-^13^C4-OncoFAP-NODAGA, or [^nat^F]AlF-^13^C4-OncoFAP-NOTA) were added to the mixture. Samples were then homogenized with a tissue lyser (TissueLyser II, QIAGEN) for 15 minutes at 30 Hz. After homogenization, samples were centrifugated at 15’000 g for 10 minutes and supernatants were dried at room temperature with a *vacuum* centrifuge (Eppendorf). Pellets were then resuspended in 1 mL solution containing 3% ACN and 0.1% of TFA and subsequently purified on Oasis HLB SPE columns (Waters) following instructions indicated by the manufacturer. Eluates were dried under *vacuum* at room temperature, resuspended in 400 μL 3% ACN and 0.1% of TFA and further purified on Macrospin column (Harvard Apparatus). Eluates were then dried under vacuum at room temperature.

Dried samples were finally resuspended in 30 μL of a solution containing 3% of ACN and 0.1% of FA. Each sample (1.5 μL, 5% of the total) was then injected in the nanoLC-HRMS system.

### nanoLC-HRMS analysis

Chromatographic separation was carried out on an Acclaim PepMap RSLC column (50 μm x 15 cm, particle size 2 μm, pore size 100 Å, Thermo Fisher Scientific) with a gradient program from 95% A (H_2_O, 0.1% FA), 5% B (ACN 0.1% FA) to 5% A, 95% B in 45 minutes on an Easy nanoLC 1000 (Thermo Fisher Scientific). The LC system was coupled to a Q-Exactive mass spectrometer (Thermo Fisher Scientific) via a Nano Flex ion source (Thermo Fisher Scientific). Ionization was carried out with 2 kV of spray voltage, 250 °C of capillary temperature, 60 S-lens RF level. The mass spectrometer was operating in Single Ion Monitoring (SIM) mode, following the mass range reported in **Table 1**. The detector was working in positive ion mode with the following parameters: resolution 70000 (FWHM at 200 m/z), AGC target 5 x 10^4^, and maximum injection time 200 ms.

**Table 1:**
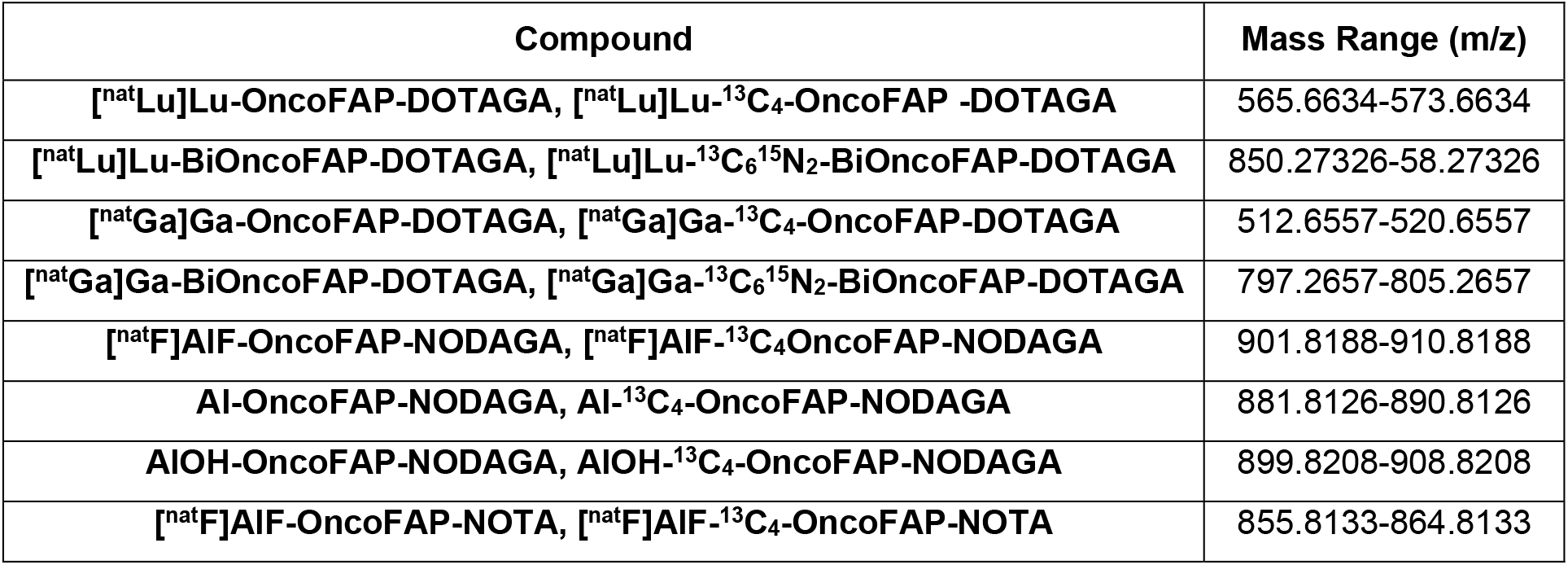
mass range windows for the SIM mode of the mass spectrometer.

| Compound | Mass Range (m/z) |
| --- | --- |
| [<sup>nat</sup>Lu]Lu-OncoFAP-DOTAGA, [<sup>nat</sup>Lu]Lu-<sup>13</sup>C<sub>4</sub>-OncoFAP -DOTAGA | 565.6634-573.6634 |
| [<sup>nat</sup>Lu]Lu-BiOncoFAP-DOTAGA, [<sup>nat</sup>Lu]Lu-<sup>13</sup>C<sub>6</sub><sup>15</sup>N<sub>2</sub>-BiOncoFAP-DOTAGA | 850.27326-58.27326 |
| [<sup>nat</sup>Ga]Ga-OncoFAP-DOTAGA, [<sup>nat</sup>Ga]Ga-<sup>13</sup>C<sub>4</sub>-OncoFAP-DOTAGA | 512.6557-520.6557 |
| [<sup>nat</sup>Ga]Ga-BiOncoFAP-DOTAGA, [<sup>nat</sup>Ga]Ga-<sup>13</sup>C<sub>6</sub><sup>15</sup>N<sub>2</sub>-BiOncoFAP-DOTAGA | 797.2657-805.2657 |
| [<sup>nat</sup>F]AlF-OncoFAP-NODAGA, [<sup>nat</sup>F]AlF-<sup>13</sup>C<sub>4</sub>OncoFAP-NODAGA | 901.8188-910.8188 |
| Al-OncoFAP-NODAGA, Al-<sup>13</sup>C<sub>4</sub>-OncoFAP-NODAGA | 881.8126-890.8126 |
| AlOH-OncoFAP-NODAGA, AlOH-<sup>13</sup>C<sub>4</sub>-OncoFAP-NODAGA | 899.8208-908.8208 |
| [<sup>nat</sup>F]AlF-OncoFAP-NOTA, [<sup>nat</sup>F]AlF-<sup>13</sup>C<sub>4</sub>-OncoFAP-NOTA | 855.8133-864.8133 |

### Data analysis

Peak areas of analytes and internal standards were integrated, and corresponding ratios calculated. Ratios were then transformed in pmol/g of wet tissue using single concentration external calibration points (**Table S1**) and corrected by the total weight of the sample analyzed. The percentage of injected dose per gram (%ID/g) was finally determined by normalizing the value based on the total dose injected in the mouse (i.e., 5 nmol, or 10 nmol). All biodistribution experiments were performed using 3 mice per experimental condition. Graphs express mean ± standard deviation values. Multiple t-test was done to compare %ID/g in each organ for i) [^nat^Lu]Lu-OncoFAP-DOTAGA and [^177^Lu]Lu-OncoFAP-DOTAGA; ii) [^nat^Lu]Lu-BiOncoFAP-DOTAGA and [^177^Lu]Lu-BiOncoFAP-DOTAGA; iii) [^nat^Ga]Ga-OncoFAP-DOTAGA and [^nat^Lu]Lu-OncoFAP-DOTAGA; iiii) [^nat^Ga]Ga-BiOncoFAP-DOTAGA and [^nat^Lu]Lu-BiOncoFAP-DOTAGA. Data analysis was performed with Thermo Xcalibur Qual Broswer v2.2 (Thermo Fisher Scientific) and Prism8 (GraphPad).

## Results and Discussion

### Method Development

We have developed an innovative LC-MS method to measure SMMCs targeting FAP in *in-vivo* biodistribution experiments. **Figure 1** presents a summary of the methodology, the schematic workflow of the experimental design, and the sample preparation procedure. The method was applied for the quantification of OncoFAP-derivatives (**Figure 2**) in biological matrices. [^nat^Lu]Lu-OncoFAP-DOTAGA, [^nat^Lu]Lu-BiOncoFAP-DOTAGA, [^nat^Ga]Ga-OncoFAP-DOTAGA, [^nat^Ga]Ga-BiOncoFAP-DOTAGA, [^nat^F]AlF-OncoFAP-NODAGA, and [^nat^F]AlF-OncoFAP-NOTA were obtained in high purity and yields following well-established and previously described procedures^11^ (**Supplementary Info**).

**Figure 1:**
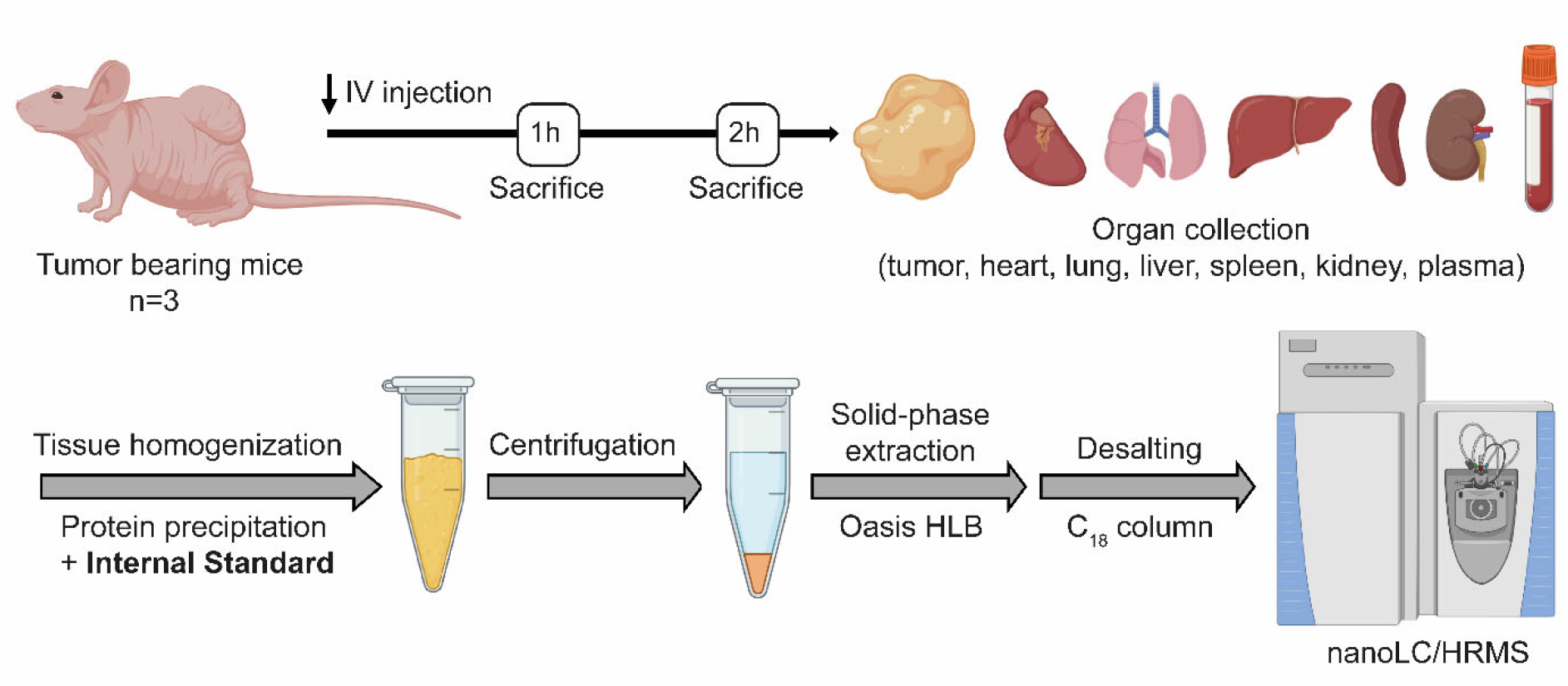
Schematic workflow showing the experimental design and the sample preparation procedure. Mice (n=3) were injected intravenously with different FAP-targeting compounds (250 nmol/Kg or 500 nmol/Kg). Mice were sacrificed 1 or 2 hours post-injections, tissues were harvested and biospecimens processed and analyzed by nanoLC-HRMS.

**Figure 2:**
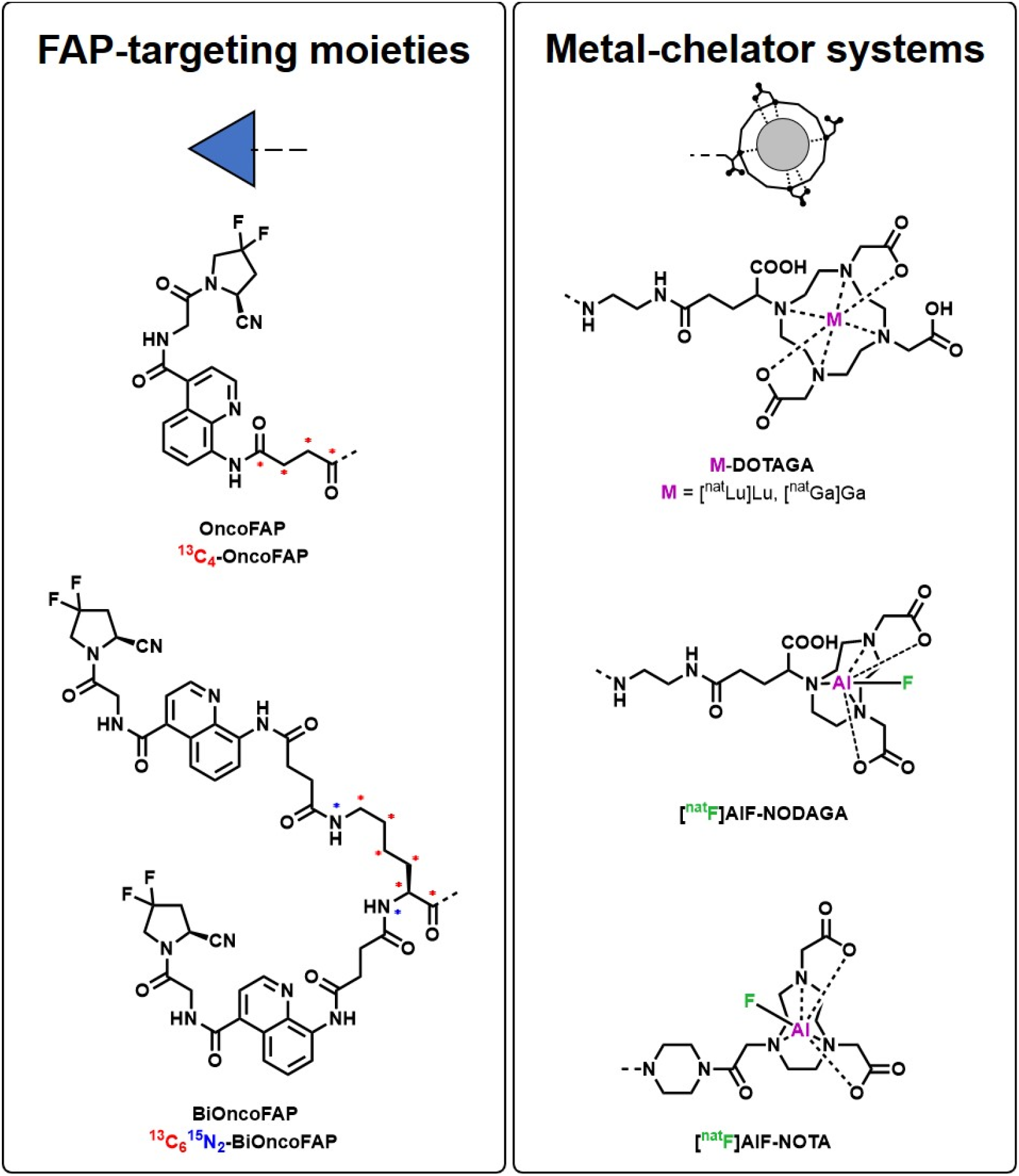
Chemical structures of targeting moieties (i.e., OncoFAP, and BiOncoFAP), and structures of metal chelators (i.e., DOTAGA, NODAGA, and NOTA) used in this study. Colored asterisks indicate the position of either ^13^C or ^15^N incorporated in the internal standard.

Ionization efficiency and chromatography parameters (e.g., peak shape, resolution, and retention time) were evaluated for each compound by injecting an aqueous solution in the LC-MS system and method parameters were tuned accordingly. Sample preparation was developed with the purpose to maximize the enrichment of the analytes after extraction from biological matrices prior to the quantification by LC-MS (**Figure 1**). Matrix effect and recovery after sample preparation were evaluated by spiking known amounts of the analytes in SK-RC-52 or HT-1080 tumor tissues from untreated animals (“blank” tumors) (**Table S2**). Matrix effect was found to be variable between different analytes, with a ~90% loss of signal for DOTAGA-conjugates and ~50% for NOTA and NODAGA derivatives. A recovery of ~50% was instead observed for all analyzed compounds.

Due to significant loss of signal observed and to obtain a reliable quantification, we decided to include highly pure and stable isotopically labeled internal standards (ISs) before sample preparation. ISs were synthesized following identical synthetic routes utilized for the analytes, replacing key reagents with corresponding ^13^C or ^15^N labeled compounds. ^13^C4 succinic anhydride was used as building block for OncoFAP-based molecules, while ^13^C6^15^N_2-L_-Lysine was used for BiOncoFAP derivatives (**Figure 2**). Sensitivity of the analysis was further improved by operating the MS instrument in SIM mode. The simultaneous detection of analytes and of corresponding ISs in the same scan range allowed to minimize the interference caused by co-elution of contaminants. The increased sensitivity of the SIM mode and the use of isotopically labelled ISs allowed us to detect and precisely measure the analytes in all tissues analyzed despite the not complete recovery after sample preparation, and the high matrix effects for some derivatives.

### Biodistribution of [^nat^Lu]Lu-OncoFAP-DOTAGA and [^nat^Lu]Lu-BiOncoFAP-DOTAGA

We have recently presented biodistribution results obtained with [^177^Lu]Lu**-**OncoFAP-DOTAGA and [^177^Lu]Lu-BiOncoFAP-DOTAGA in HT-1080.hFAP tumor bearing mice that show favorable and selective accumulation of both compounds in FAP-positive tumors^11^. With the aim of benchmarking our LC-MS methodology with radioactivity-based results obtained with [^177^Lu]Lu-OncoFAP-DOTAGA and [^177^Lu]Lu-BiOncoFAP-DOTAGA, we studied quantitative biodistributions of their cold counterparts. Mice bearing HT-1080.hFAP tumors were treated with [^nat^Lu]Lu-OncoFAP-DOTAGA and [^nat^Lu]Lu-BiOncoFAP-DOTAGA and sacrificed 1 hour after injection. Sensitivity of LC-MS analysis was high enough to accurately measure both compounds in any specimen analyzed. As expected, both molecules accumulated selectively in FAP-positive tumors. The BiOncoFAP product exhibited a higher tumor uptake as compared to its monovalent counterpart (**Figures 3, Table S3**). Uptake values in tumor and healthy organs (in %ID/g) measured by the LC-MS methodology were comparable to values obtained with radioactivity-based quantification^11^. No significant differences were observed between the two analytical techniques (p>0.05 multiple t test, **Table S4**) (**Figures 3**). Orthogonal validation further confirms the accuracy of the newly developed MS quantification method, thereby opening new opportunities for non-radioactive LC-MS analysis of SMMCs.

**Figure 3:**
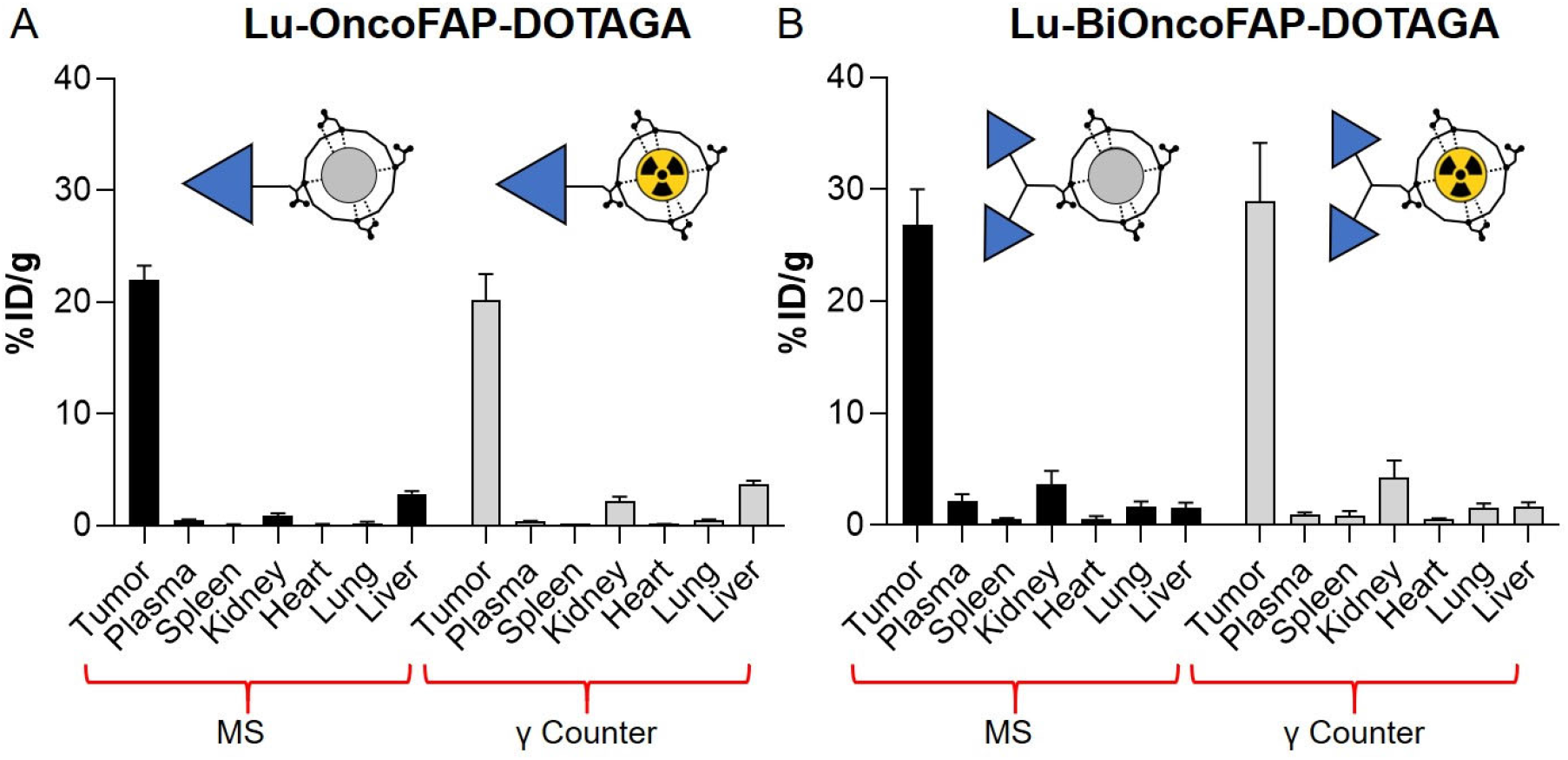
Comparison of MS and γ Counter^11^ biodistribution data for A) Lu-OncoFAP-DOTAGA and B) Lu-BiOncoFAP-DOTAGA. Mice bearing HT-1080.hFAP tumors were sacrificed 1 h after intravenous administration (250 nmol/kg). No significant differences were observed between biodistribution results measured by MS and by radioactivity (p > 0.05, multiple t test, **table S4**).

### Biodistribution of [^nat^Ga]Ga-OncoFAP-DOTAGA and [^nat^Ga]Ga-OncoFAP-DOTAGA

Encouraged by these results, we further applied our methodology for the assessment of the biodistributions of [^nat^Ga]Ga-OncoFAP-DOTAGA and [^nat^Ga]Ga-BiOncoFAP-DOTAGA, cold molecular counterparts of two novel diagnostic radiotracers (i.e., [^68^Ga]Ga-OncoFAP-DOTAGA and [^68^Ga]Ga-BiOncoFAP-DOTAGA)^6^ (**Figure 4, table S5**). Similarly to Lutetium derivatives, both compounds showed a consistent tumor uptake and no significant uptake in healthy organs, with favorable tumor-to-organ ratios. Uptake values obtained for [^nat^Ga]Ga-OncoFAP-DOTAGA and [^nat^Ga]Ga-BiOncoFAP-DOTAGA showed no significant differences with [^nat^Lu] derivatives (p>0.05 multiple t test, **table S4**), except for a lower liver uptake observed for both ^nat^Ga counterparts (p<0.05 multiple t test **table S4**). Despite some minor differences, probably caused by the intrinsic physical-chemical properties of the molecules, the tumor accumulations of the mono and bivalent OncoFAP-metal-conjugates were remarkable in every biodistribution experiment performed. Our results further confirm the versatility of this class of molecules as targeted radiopharmaceuticals^6,10,11^.

**Figure 4:**
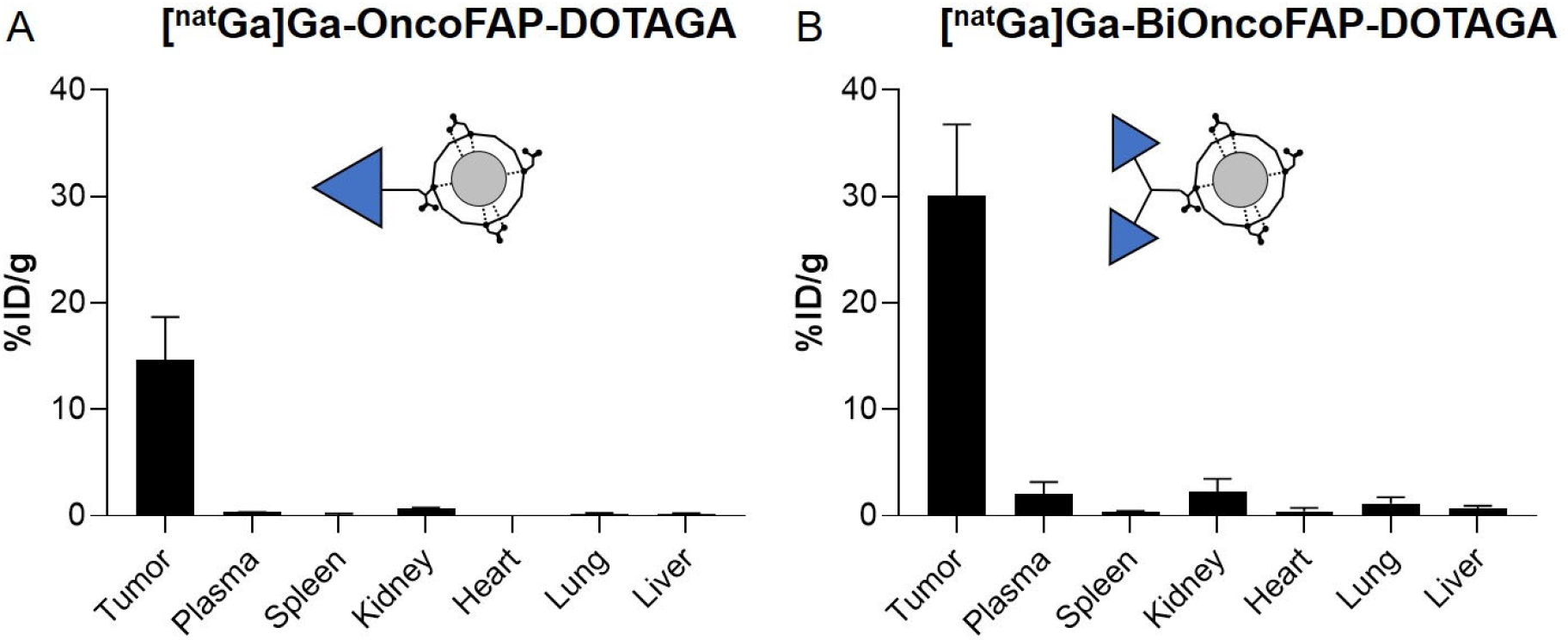
MS biodistribution results of A) [^nat^Ga]Ga-OncoFAP-DOTAGA and B) [^nat^Ga]Ga-BiOncoFAP-DOTAGA. Mice bearing HT-1080.hFAP tumors were sacrificed 1 h after intravenous administration (250 nmol/kg).

### Biodistribution of [^nat^F]AlF-OncoFAP-NODAGA and [^nat^F]AlF-OncoFAP-NOTA

Besides Gallium-68 and Lutetium-177, Fluorine-18 is also frequently used for nuclear medicine applications in cancer patients^17–19^. Fluorine-18 can be covalently bound to tumor targeting agents (i.e., [^18^F]-Fluorodeoxyglucose)^20^ or indirectly incorporated as counter-ion in Aluminum complexes^19^. Among several metal chelators commercially available for chelating Aluminum Fluoride, the most widely used are NODAGA and NOTA^19,21,22^. For this purpose, OncoFAP-NODAGA and OncoFAP-NOTA were synthetized and labeled with [^nat^F]AlF_3_. After conjugation three peaks corresponding to three different species of [^nat^F]AlF-OncoFAP-NODAGA were identified by MS (**Figure 5A**), while only one peak was observed for [^nat^F]AlF-OncoFAP-NOTA (**Figure 5B**). The three peaks of [^nat^F]AlF-OncoFAP-NODAGA could be unambiguously assigned to AlOH-OncoFAP-NODAGA (Rt = 13.56 min), Al-OncoFAP-NODAGA (Rt = 15.34 min) and [^nat^F]AlF-OncoFAP-NODAGA (Rt = 16.42 min) (**Figure 5A**). Among the three species present, the most abundant one corresponds to Al-OncoFAP-NODAGA. This observation is not unexpected since it is known that NODAGA N3O3 configuration is not favorable for AlF_3_ chelation^18,19^. By observing the structures of NODAGA and NOTA chelators, it is possible to note the presence of an extra carboxylic group in the NODAGA structures that saturates Aluminum coordination sphere^18,22^, resulting in the loss of the Fluoride anion (**Figure S1**). On the contrary, NOTA N3O2 configuration, facilitates the AlF_3_ chelation not having a carboxylic group competing with the Fluoride interaction^18,23^ (**Figure S1**). Biodistribution experiments were conducted in SK-RC-52.hFAP bearing mice treated with either [^nat^F]AlF-OncoFAP-NODAGA or [^nat^F]AlF-OncoFAP-NOTA (**Figure 5C-D**), following the same technical procedures and quantification methods described above. The %ID/g of OncoFAP-NODAGA was calculated as sum of the three different molecular species since it was not possible to separate them after cold-labeling and therefore a mixture of the three was injected in the animals. Our newly developed LC-MS methodology allowed us to precisely follow the biodistribution of each single molecular species of OncoFAP-NODAGA, thereby resulting into a more accurate analysis than classical radioactive-based method which are limited to the [^18^F]AlF-OncoFAP-NODAGA only species (**Figure S2, Table S6**). Notably, both molecules are characterized by favorable tumor-to-organ ratios and do not show any significant uptake in healthy organs. [^nat^F]AlF-OncoFAP-NOTA shows a better performance since it exhibited a higher tumor uptake compared to OncoFAP-NODAGA (i.e., 12.83±0.87 vs 5.19±1.55 %ID/g.) (**Figure 5C-D. Table S7**).

**Figure 5:**
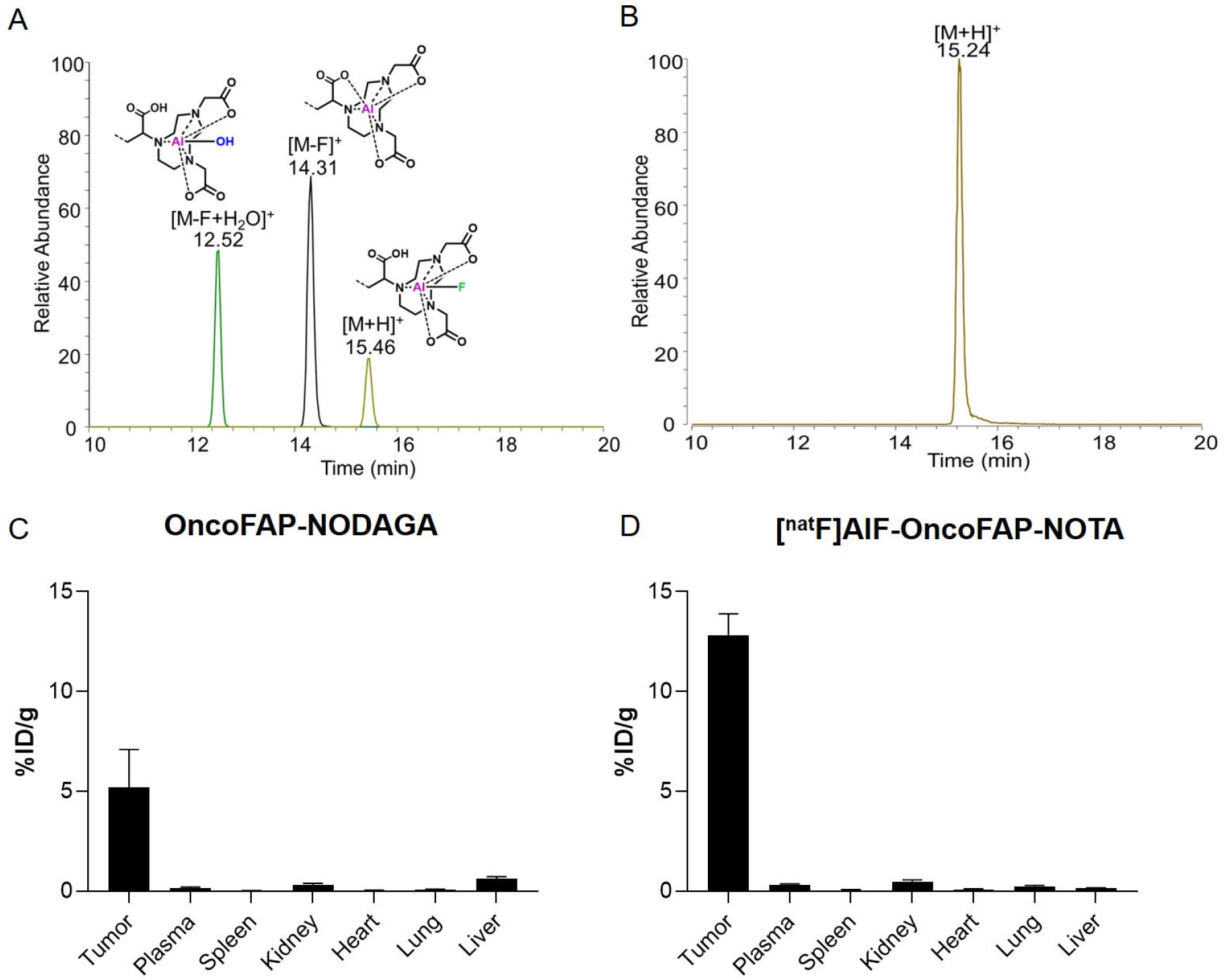
Chromatograms of A) OncoFAP-NODAGA and B) OncoFAP-NOTA, after labeling with Al^nat^F; MS biodistribution results of C) OncoFAP-NODAGA and D) [^nat^F]AlF-OncoFAP-NOTA. Chromatogram (A) evidences the presence of three different molecular species eluting at different retention times: AlOH-OncoFAP-NODAGA (Rt = 13.56 min), Al-OncoFAP-NODAGA (Rt = 15.34 min) and [^nat^F]AlF-OncoFAP DOTAGA (Rt = 16.42 min); The chromatogram of OncoFAP-NOTA (B) shows the presence of one single peak corresponding to [^nat^F]F-OncoFAP-NOTA (Rt = 15.24 min).For biodistribution experiments (C-D) mice bearing SK-RC-52.hFAP tumors were sacrificed 2 h after intravenous administration (500 nmol/kg). For OncoFAP-NODAGA (C), the %ID/g was calculated as sum of the three different molecular species (AlOH-OncoFAP-NODAGA, Al-OncoFAP-NODAGA and [^nat^F]AlF-OncoFAP DOTAGA) observed and measured by MS. For OncoFAP-NOTA (D) the %ID/g refers to the only molecular species ([^nat^F]F-OncoFAP-NOTA) measured by MS.

## Conclusion

In this article, we have reported the development and the application of a novel quantitative LC-MS methodology for the assessment of the *in vivo* biodistribution analysis of tumor targeting SMMCs. The method presented here is reliable and accurate in providing biodistribution data which are fully comparable to those obtained by radioactive-based experiments. The use of stable isotopes and the successful quantification of SMMCs by MS represent a valid alternative to classical approaches, opening new possibilities in the discovery of novel radiopharmaceuticals. Quantification methodologies based on radioactivity are often used and can be very accurate, if a suitable radiolabeling procedure is used. However, regulatory constraints (e.g., the need for dedicated laboratories and infrastructure) as well as safety concerns (e.g., exposure of the operator to harmful radiations) limit the applicability of radioactivity-based methodologies, both in academia and in industry. In this context, our newly developed LC-MS methodology represents a valuable orthogonal, safe, green, and easy-to-implement alternative. Our vision considers further integration of LC-MS quantification in drug development of targeted SMRCs. We envisage a further expansion of the technology described in this article for the assessment of *in vivo* biodistribution of other classes of targeted drugs (e.g., Small Molecule-Drug Conjugates and antibody-cytokine fusion proteins). Overall, the reduction of the radioactive or cytotoxic compounds during discovery phases will translate into an enhanced throughput and, ultimately, into higher possibilities of identification of novel clinical candidates.

## Abbreviations

ACN: Acetonitrile
DOTAGA: 2-(4,7,10-tris(carboxymethyl)-1,4,7,10-tetraazacyclododecan-1-yl)pentanedioic acid
EDTA: Ethylenediaminetetraacetic acid
FA: Formic acid
FAP: Fibroblast activation protein
HPLC: High pressure liquid chromatography
HRMS: High resolution mass spectrometry
ICP-MS: Inductively coupled plasma mass spectrometry
ID/g: Injected dose per gram
IS: Internal standard
LC: Liquid chromatography
LC-MS: Liquid chromatography – Mass spectrometry
MS: Mass spectrometry
NODAGA: 2-(4,7-bis(carboxymethyl)-1,4,7-triazonan-1-yl)pentanedioic acid
NOTA: 2,2’-(7-carboxy-1,4,7-triazonane-1,4-diyl)diacetic acid
PBS: Phosphate-buffered saline
SIM: Single ion monitoring
SMMC: Small molecule metal conjugate
SMRC: Small molecule radio conjugate
SPE: Solid phase extraction
TFA: Trifluoroacetic acid

