## Supplementary Information for "A mass spectrometry-based method for the determination of *in vivo* biodistribution of tumor targeting small molecule-metal conjugates"

### Index

### Chemical synthesis

All reagents and solvents were purchased from Sigma Aldrich, VWR, Combi-Blocks, CheMatech and used as supplied.

High Pressure Liquid Chromatography/Mass-Spectrometry (HPLC/MS) spectra presented were recorded on an Agilent 6100 Series Single Quadrupole MS system combined with an Agilent 1200 Series LC, using an InfinityLab Poroshell 120 EC-C18 Column (4.6 × 50 mm, 2.7 µm particle size, 120 Å pore size) at a flow rate of 0.8 mL/min.

| Time (min) | Water + 0.1% FA | ACN + 0.1% FA | Flow (mL/min) |
|---|---|---|---|
| 0.00 | 90.0% | 10% | 0.800 |
| 0.50 | 90.0% | 10% | 0.800 |
| 3.00 | 0.0% | 100% | 0.800 |
| 3.50 | 0.0% | 100% | 0.800 |

Reversed-phase high-pressure liquid chromatography (RP-HPLC) purifications were performed on an Agilent 1200 Series RP-HPLC with PDA UV detector, using a Synergi MAX-RP C18 column (10 × 250 mm, 10 µm particle size, 80 Å pore size) at a flow rate of 5 mL/min with linear gradients of solvents A and B (A = Millipore water with 0.1% TFA, B = ACN with 0.1% TFA).

Reverse-phase chromatography purifications were performed on a Büchi Sepacore using a FlashPure ID C18 irregular column (24g, particle size 40 µm, pore size: 53-80 Å) at a flow rate of 30 mL/min with linear gradients of solvents A and B (A = Millipore water with 0.1% FA, B = ACN with 0.1% FA).

Direct phase column chromatography purifications were performed on a CombiFlash NextGen 300+ using RediSep Silver normal phase silica irregular columns (particle size: 40–63 µm, mesh size: 230–400, pore size: 60 Å) with linear gradients of organic solvents.

High-Resolution mass spectrometry (HR-MS) were performed on a Q Exactive Mass Spectrometer (Thermo Fisher Scientific). The analyte was injected directly into the MS at a flow rate of 4 µL/min. Spectra were obtained with a resolution of 70000. OncoFAP-DOTAGA, [$^{nat}$Lu]Lu-OncoFAP-DOTAGA, [$^{nat}$Ga]Ga-OncoFAP-DOTAGA, BiOncoFAP-DOTAGA, [$^{nat}$Lu]Lu-BiOncoFAP-DOTAGA, were synthetized according to literature procedures[1–3]

### Synthetic schemes

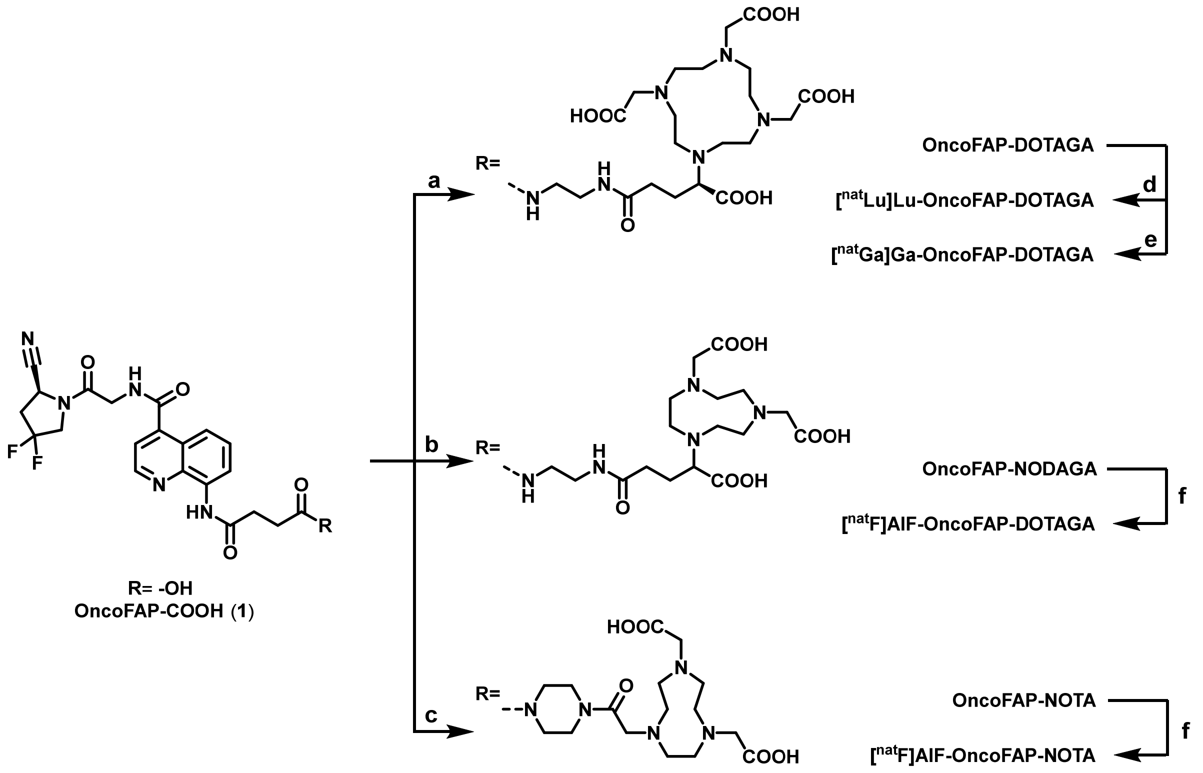

**Scheme S1.** Synthetic routes for the synthesis of OncoFAP-DOTAGA, [$^{nat}$Lu]Lu-OncoFAP-DOTAGA, [$^{nat}$Ga]Ga-OncoFAP-DOTAGA, OncoFAP-NODAGA, [$^{nat}$F]AlF-OncoFAP-NODAGA, OncoFAP-NOTA and [$^{nat}$F]AlF-OncoFAP-NOTA. Reagents and conditions: a) (*i*) *N*-hydroxysuccinimide, HATU, DIPEA, dry DMSO, r.t., 30 min, (*ii*) (*R*)-DOTAGA-$NH_2$, water, r.t. o.n.; b) (*i*) *N*-hydroxysuccinimide, HATU, DIPEA, DMF, r.t., 30 min, (*ii*) NODAGA-$NH_2$, water, r.t. o.n.; c) (*i*) *N*-Boc-piperazine, HATU, DIPEA, r.t., 30 min, (*ii*) TFA/DCM 30% v/v, r.t., 3h, (*iii*) NOTA-NHS, DIPEA, DMF, r.t., 1h; d) $LuCl_3$ * $6H_2O$, acetate buffer pH 8, 0.05 N HCl, DMSO, 95°C, 15 min; e) $GaCl_3$, acetate buffer pH 4.5, DMSO, 95°C, 15 min; f) $AlF_3$, acetate buffer pH 4, DMSO, 95°C, 15 min.

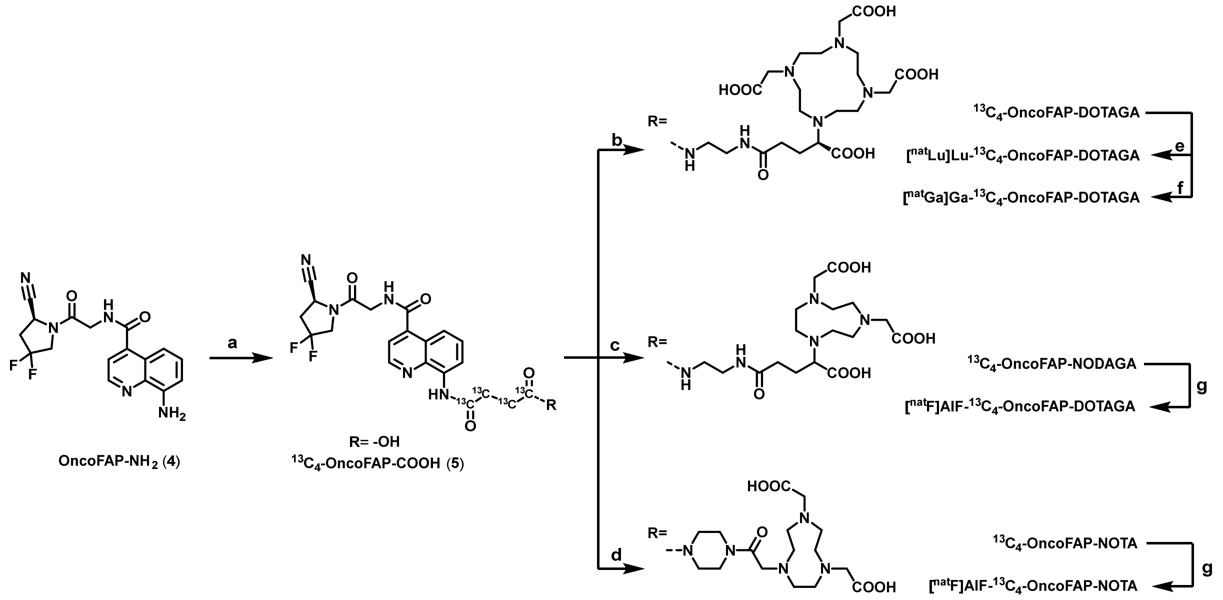

**Scheme S2.** Synthetic routes for the synthesis of $^{13}C_4$-OncoFAP-DOTAGA, [natLu]Lu-$^{13}C_4$-OncoFAP-DOTAGA, [natGa]Ga-$^{13}C_4$-OncoFAP-DOTAGA, $^{13}C_4$-OncoFAP-NODAGA (X), [natF]AlF-$^{13}C_4$-OncoFAP-NODAGA,$^{13}C_4$-OncoFAP-NOTA and [natF]AlF-$^{13}C_4$-OncoFAP-NOTA (X).Reagents and conditions: a) $^{13}C_4$-succinic anhydride, DMAP, THF, 55°C, 1h; b) (*i*) *N*-hydroxysuccinimide, HATU, DIPEA, dry DMSO, r.t., 30 min, (*ii*) (*R*)-DOTAGA-$NH_2$, water, r.t. o.n.; c) (*i*) *N*-hydroxysuccinimide, HATU, DIPEA, DMF, r.t., 30 min, (*ii*) NODAGA-$NH_2$, water, r.t. o.n.; d) (*i*) *N*-Boc-piperazine, HATU, DIPEA, r.t., 30 min, (*ii*) TFA/DCM 30% v/v, r.t., 3h, (*iii*) NOTA-NHS, DIPEA, DMF, r.t., 1h; e) $LuCl_3$ * $6H_2O$, acetate buffer pH 8, 0.05 N HCl, DMSO, 95°C, 15 min; f) $GaCl_3$, acetate buffer pH 4.5, DMSO, 95°C, 15 min; g) $AlF_3$, acetate buffer pH 4, DMSO, 95°C, 15 min.

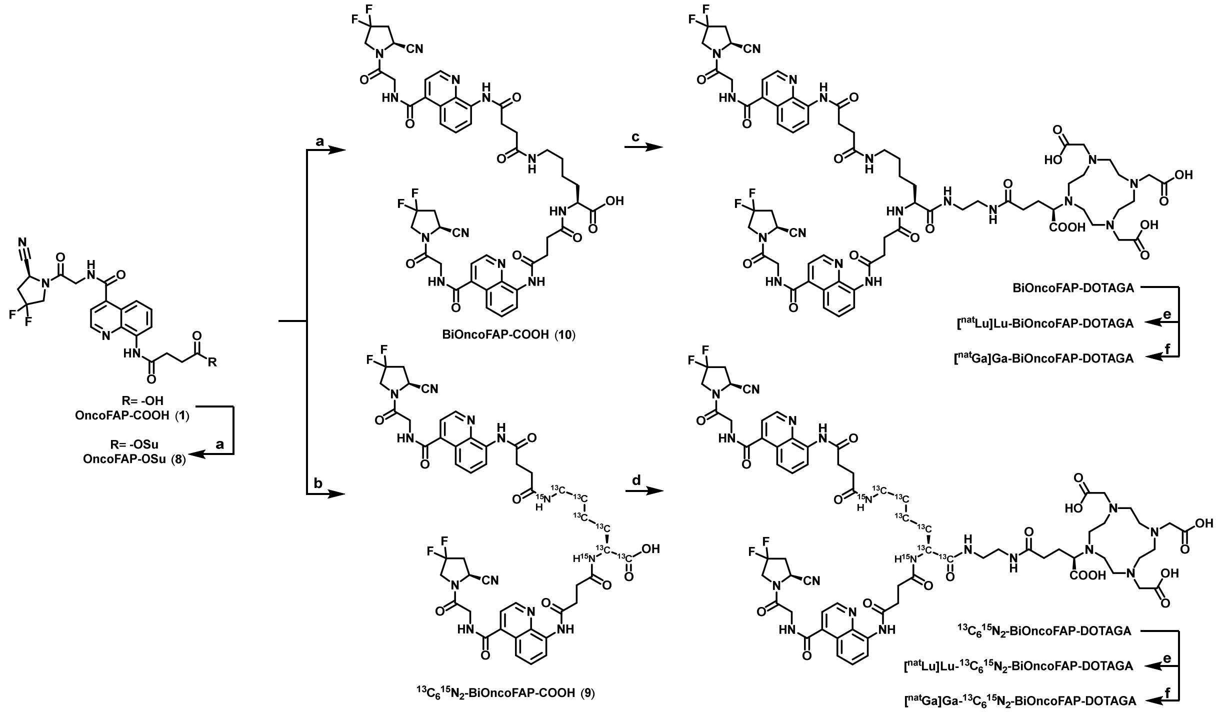

**Scheme S3.** Synthetic routes for the synthesis of BiOncoFAP-DOTAGA, [$^{nat}$Lu]Lu-BiOncoFAP-DOTAGA, [$^{nat}$Ga]Ga-BiOncoFAP-DOTAGA, $^{13}C_6{}^{15}N_2$-BiOncoFAP-DOTAGA, [$^{nat}$Lu]Lu-$^{13}C_6{}^{15}N_2$BiOncoFAP-DOTAGA and [$^{nat}$Ga]Ga-$^{13}C_6{}^{15}N_2$BiOncoFAP-DOTAGA. Reagents and conditions: a) L-Lysine, DIPEA, DMSO, r.t., o.n.; b) $^{13}C_6{}^{15}N_2$- L-Lysine, DIPEA, DMSO, r.t., o.n.; c) (*i*) *N*-hydroxysuccinimide, HATU, DIPEA, DMF, r.t., 30 min, (*ii*) (*R*)-DOTAGA-$NH_2$, water, r.t. o.n.; e) $LuCl_3$ * $6H_2O$, acetate buffer pH 8, 0.05 N HCl, DMSO, 95°C, 15 min; f) $GaCl_3$, acetate buffer pH 4.5, DMSO, 95°C, 15 min;

**Experimental procedures**

Synthesis of ***tert*****-butyl (*****S*****)-4-(4-((4-((2-(2-cyano-4,4-difluoropyrrolidin-1-yl)-2-oxoethyl)carbamoyl)quinolin-8-yl)amino)-4-oxobutanoyl)piperazine-1-carboxylate** (**2**)

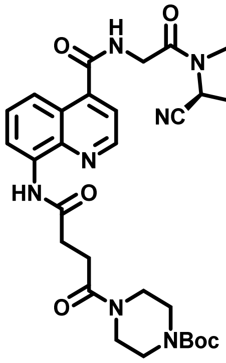

**OncoFAP-COOH** (**1**) (50 mg, 0.11 mmol, 1 eq), *N*-Boc-piperazine (24 mg, 0.13 mmol, 1.2 eq) and HATU (49 mg, 0.13 mmol, 1.2 eq) were dissolved in DMF (1 mL). DIPEA (0.08 mL, 0.44 mmol, 4 eq) was added dropwise and the mixture was stirred for 30 min at room temperature. The mixture was purified via RP flash chromatography (Büchi Sepacore equipped with FlashPure ID C18 irregular column) using a gradient of water + 0.1% FA/ACN + 0.1% FA 98:2 to 0:100 in 40 min. The desired fractions were collected and lyophilized to afford a white solid. (40 mg, 58%)

MS (ESI+) m/z 627.9 [M+H]

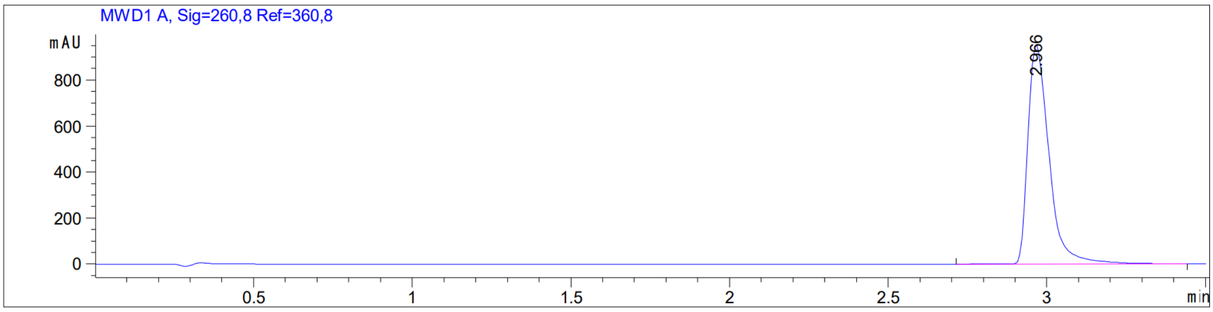

Synthesis of **(*S*)-*N*-(2-(2-cyano-4,4-difluoropyrrolidin-1-yl)-2-oxoethyl)-8-(4-oxo-4-(piperazin-1-yl)butanamido)quinoline-4-carboxamide** (3)

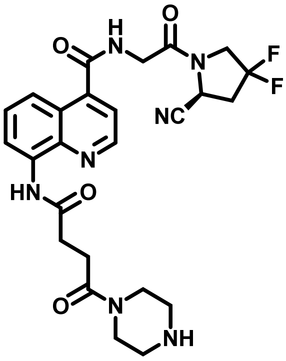

Compound (**2**) (40 mg, 0,06 mmol, 1 eq.) was dissolved in DCM (1 mL) and TFA (0.5 mL) was added dropwise. The mixture was stirred at room temperature for 3h then solvent was evaporated, and the crude was purified via RP flash chromatography (Büchi Sepacore equipped with FlashPure ID C18 irregular column) using a gradient of water + 0.1% FA/ACN + 0.1% FA 98:2 to 0:100 in 40 min. The desired fractions were collected and lyophilized to afford a colorless oil. (17 mg, 51%)

MS (ESI+) m/z 527.9 $[M+H]^+$

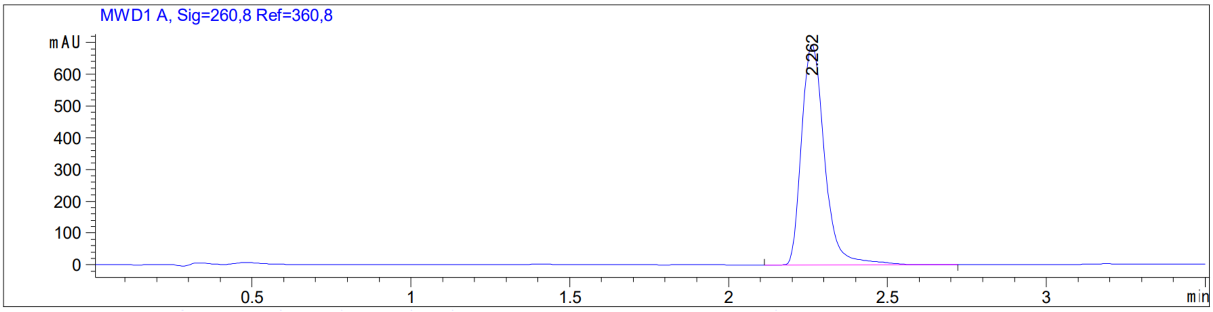

Synthesis of **(*S*)-2,2'-(7-(2-(4-(4-((4-((2-(2-cyano-4,4-difluoropyrrolidin-1-yl)-2-oxoethyl)carbamoyl)quinolin-8-yl)amino)-4-oxobutanoyl)piperazin-1-yl)-2-oxoethyl)-1,4,7-triazonane-1,4-diyl)diacetic acid, OncoFAP-NOTA**

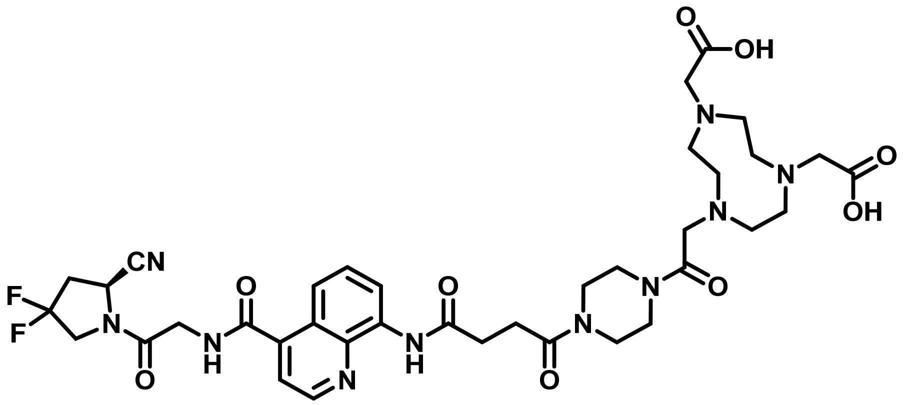

Compound (**3**) (8 mg, 0.02 mmol, 1 eq) and NOTA-NHS (19 mg, 0.04 mmol, 2 eq) were dissolved in DMF (0.2 mL). DIPEA (0.01 mL, 0.08 mmol, 4 eq) was added to the mixture and stirred for 1h at room temperature. The mixture was purified via RP-HPLC (90:10 to 0:100 water + 0.1% TFA/ACN + 0.1% TFA in 12 min). The desired fractions were collected and lyophilized to afford a white solid. (4 mg, 25%)

MS (ESI+) m/z 812.7 $[M+H]^+$

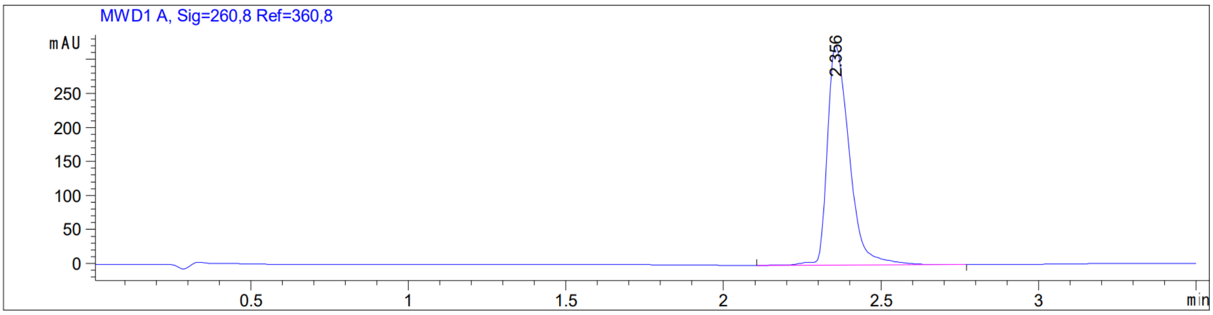

Synthesis of **OncoFAP-NODAGA**

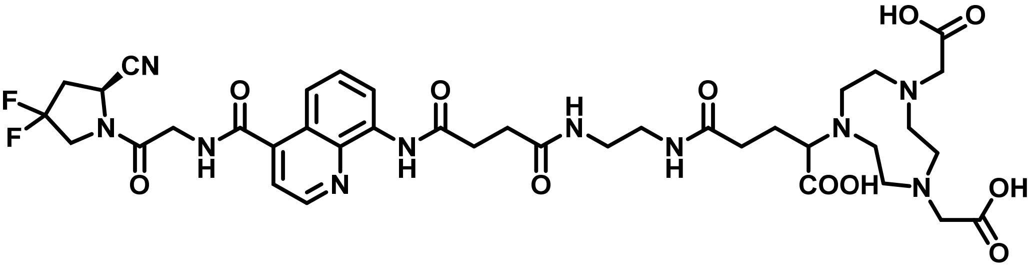

To a solution of **OncoFAP-COOH** (**1**) (50 mg, 0.11 mmol, 1 eq) in dry DMSO (1 mL) were added *N*-hydroxy succinimide (19 mg, 0.16 mmol, 1.5 eq), HATU (61 mg, 0.16 mmol, 1.5 eq) and DIPEA (57 µL, 0.44 mmol, 4 eq), the mixture at room temperature. After 30 min NODAGA-$NH_2$ (46 mg, 0.22 mmol, 2 eq) and water (0.3 mL) were added and the mixture was stirred overnight. The reaction mixture was directly purified via RP-HPLC (Agilent 1200 series system equipped with Synergi MAX-RP C18 column) using a gradient of 90:10 to 0:100 water + 0.1% TFA/ACN + 0.1% TFA in 12 min. The desired fractions were collected and lyophilized to afford a white solid. (30 mg, 32%)

MS (ESI+) m/z 859.4 $[M+H]^+$

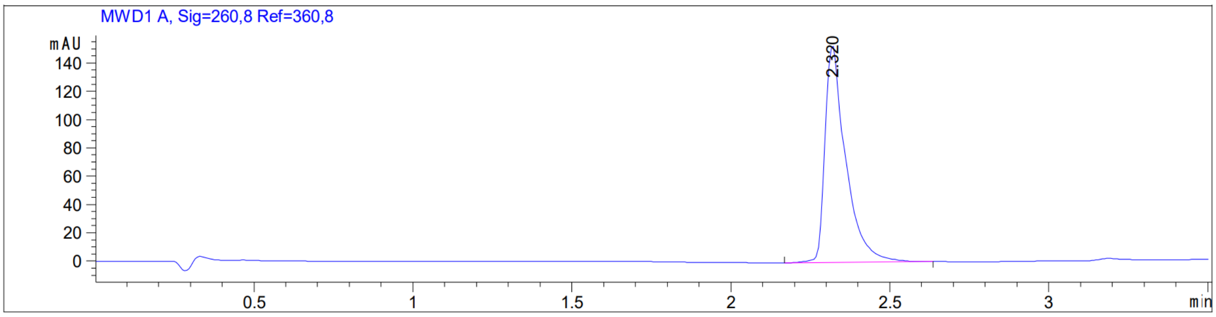

Synthesis of $^{13}C_4$-**OncoFAP-COOH** (**5**)

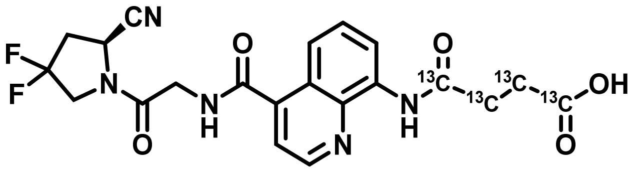

**OncoFAP-NH$_2$** (**4**) (125 mg, 0.35 mmol, 1 eq.), $^{13}C_4$-succinic anhydride (100 mg, 1.04 mmol, 3 eq.) and DMAP (8 mg, 0.07 mmol, 0.2 eq.) were dissolved in dry THF (2 mL). The mixture was heated at 55°C for 1h then cooled to room temperature, diluted with EtOAc and transferred to a separatory funnel. The mixture was washed with brine and the organic phase was dried over anhydrous $Na_2SO_4$, filtered and evaporated under reduced pressure to afford a white solid. The crude was purified via flash column chromatography (10:0 to 8:2 DCM/MeOH) to afford a white solid. (136 mg, 84%)

MS (ESI+), m/z 464.5

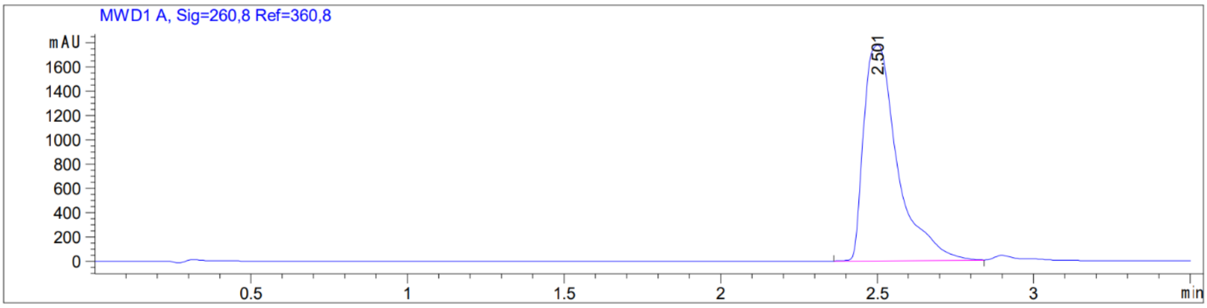

Synthesis of $^{13}$**C$_4$-OncoFAP-DOTAGA**

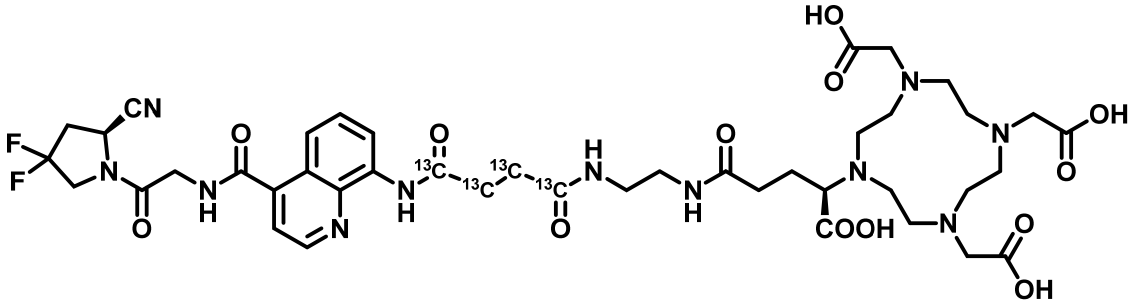

DIPEA (0.07 mL, 0.04 mmol, 4 eq.) was added dropwise to a cooled solution of **$^{13}$C$_4$-OncoFAP-COOH** (**5**) (5 mg, 0.01 mmol, 1 eq.), *N*-hydroxysuccinimide (2 mg, 0.02 mmol, 1.5 eq.) and HATU (7 mg, 0.02 mmol, 1.5 eq.). The mixture was stirred at room temperature for 30 min. Then (*R*)-DOTAGA-NH$_2$ (13 mg, 0.02 mmol, 2 eq.) was added as a solid and the suspension was stirred overnight at room temperature. The crude was diluted with water and purified via RP-HPLC (Agilent 1200 series system equipped with Synergi MAX-RP C18 column) using a gradient of 90:10 to 0:100 water + 0.1% TFA/ACN + 0.1% TFA in 16 min. Desired fractions were collected and lyophilized to afford a white solid. (8 mg, 80%)

MS (ESI+), m/z 964.4

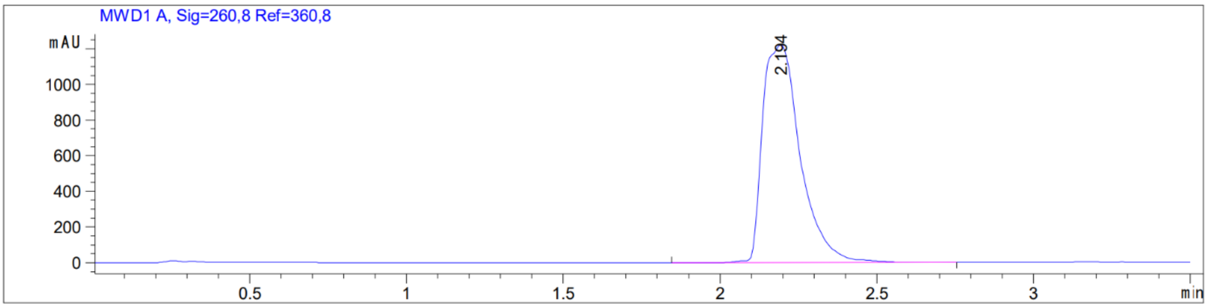

Synthesis of **[$^{nat}$Lu]Lu-$^{13}C_4$-OncoFAP-DOTAGA**

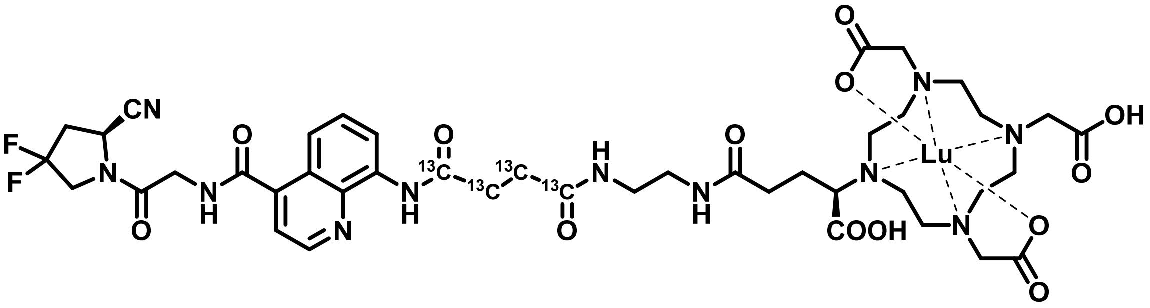

To a solution of **$^{13}C_4$-OncoFAP-DOTAGA** (1 µmol) in 300 µL acetate buffer (aqueous solution, 1 M, pH 8), a freshly prepared solution of $LuCl_3$ hexahydrate (2 eq.) in 0.05N HCl (1.5 mL) was added. The resulting mixture was stirred at 95°C for 15 minutes, then purified via RP-HPLC (Agilent 1200 series system equipped with Synergi MAX-RP C18 column) using a gradient of 90:10 to 0:100 ACN + 0.1% TFA/water + 0.1% TFA in 12 min. The desired fractions were collected and lyophilized to afford a white solid.

HRMS (ESI+) m/z 1136.33464; theoretical m/z 1136.33288

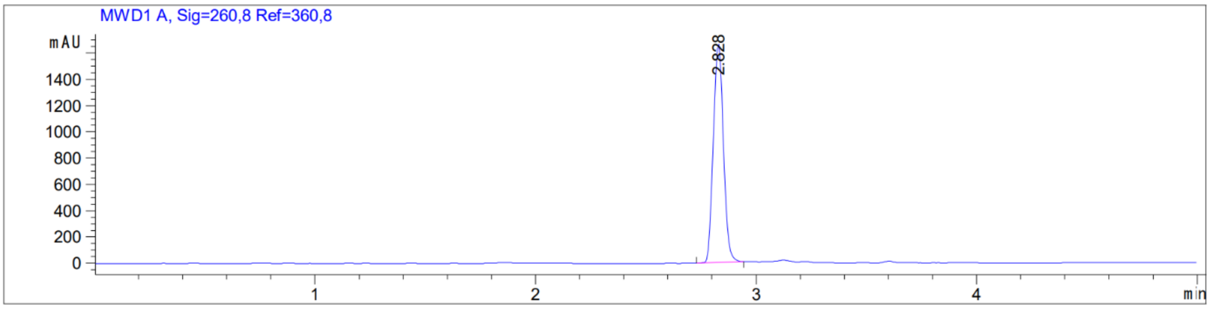

Synthesis of **[$^{nat}$Ga]Ga-$^{13}C_4$-OncoFAP-DOTAGA**

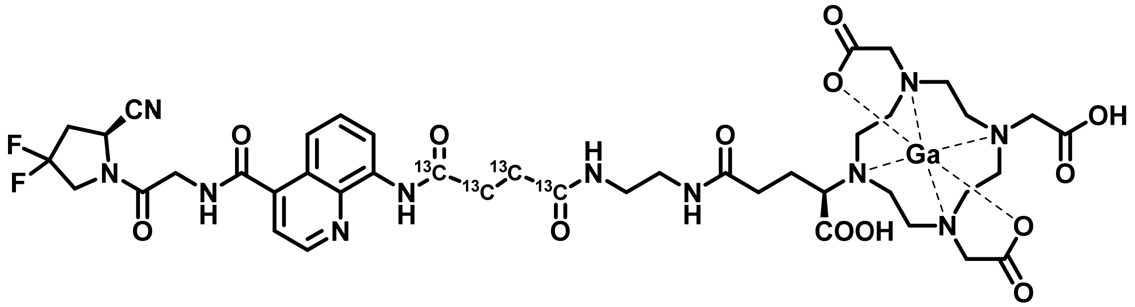

**$^{13}C_4$-OncoFAP-DOTAGA** (1 µmol) was dissolved in acetate buffer, pH=4.5 (0.1 mL). Subsequently a solution of $GaCl_3$ (10 eq.) dissolved in 1N HCl (0.01 mL) was added. The reaction was stirred at 90°C for 10 min, then cooled down to room temperature. and purified via RP-HPLC (Agilent 1200 series system equipped with Synergi MAX-RP C18 column) using a gradient of 90:10 to 0:100 water + 0.1% TFA/ACN + 0.1% TFA in 16 min. The desired fractions were collected and lyophilized to afford a pale-yellow solid.

HRMS (ESI+) m/z 1030.31929($^{69}$Ga), 1032.31928 ($^{71}$Ga); theoretical m/z 1030.31768 ($^{69}$Ga), 1032.31681 ($^{71}$Ga)

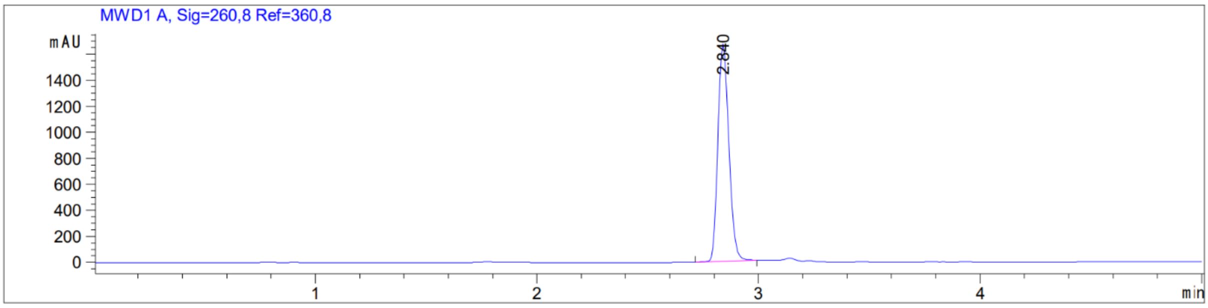

Synthesis of **$^{13}C_4$-OncoFAP-NODAGA**

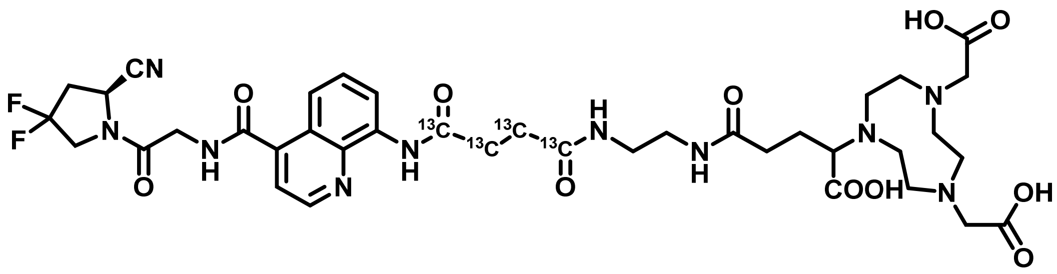

To a solution of **$^{13}C_4$-OncoFAP-COOH** (**5**) (5 mg, 0.01 mmol, 1 eq) in dry DMSO (0.1 mL) were added *N*-hydroxy succinimide (2 mg, 0.02 mmol, 1.5 eq), HATU (6 mg, 0.02 mmol, 1.5 eq) and DIPEA (6 µL, 0.04 mmol, 4 eq), the mixture at room temperature. After 30 min NODAGA-$NH_2$ (5 mg, 0.02 mmol, 2 eq) and water (0.03 mL) were added and the mixture was stirred for overnight. The reaction mixture was directly purified via RP-HPLC (Agilent 1200 series system equipped with Synergi MAX-RP C18 column) using a gradient of 90:10 to 0:100 water/ACN + 0.1% TFA in 12 min. The desired fractions were collected and lyophilized to afford a white solid. (5 mg, 56%)

MS (ESI+) m/z 863.4 $[M+H]^+$

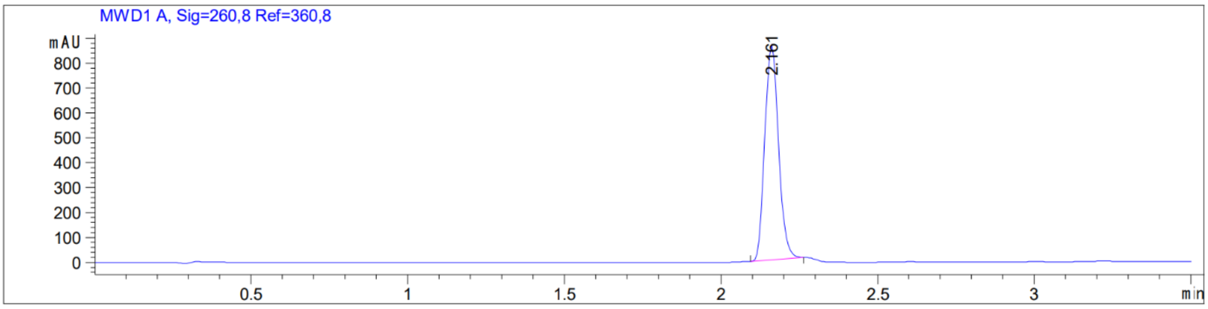

Synthesis of **[$^{nat}$F]AlF-$^{13}$C$_4$-OncoFAP-NODAGA**

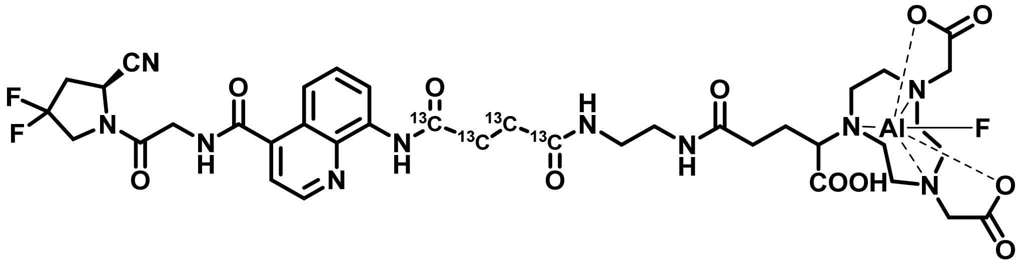

**$^{13}$C$_4$-OncoFAP-NODAGA** (1 µmol) was dissolved in a mixture of DMSO (0.05 mL) and acetate buffer pH=4 (0.2 mL). $AlF_3$ (10 eq.) was added and the mixture was heated at 95°C for 15 min. Then the mixture was purified via RP-HPLC (Agilent 1200 series system equipped with Synergi MAX-RP C18 column) using a gradient of 95:5 to 0:100 water + 0.1% TFA/ACN + 0.1% TFA in 20 min. The desired fractions were collected and lyophilized to afford a white solid.

HRMS (ESI+) m/z 907.33373; theoretical m/z = 907.33219

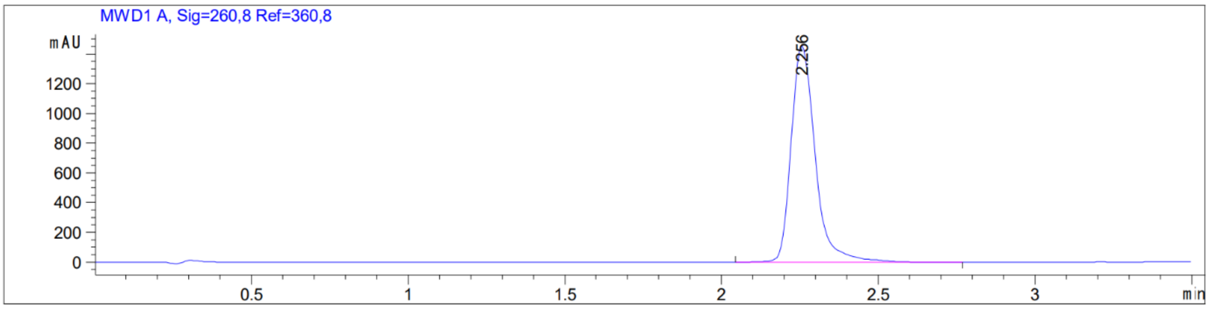

Synthesis of **$^{13}C_4$-OncoFAP-piperazine-Boc** (**6**)

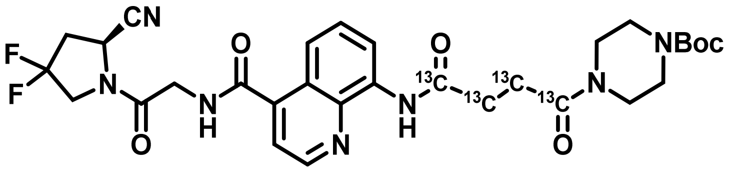

**$^{13}C_4$-OncoFAP-COOH** (**5**) (25 mg, 0.06 mmol, 1 eq), *N*-Boc-piperazine (12 mg, 0.07 mmol, 1.2 eq) and HATU (25 mg, 0.06 mmol, 1.2 eq) were dissolved in DMF (0.5 mL). DIPEA (0.04 mL, 0.22 mmol, 4 eq) was added dropwise and the mixture was stirred for 30 min at room temperature. The mixture was purified via RP flash chromatography (Büchi Sepacore equipped with FlashPure ID C18 irregular column) using a gradient of water + 0.1% FA/ACN + 0.1% FA 98:2 to 0:100 in 40 min. The desired fractions were collected and lyophilized to afford a white solid. (22 mg, 63%)

MS (ESI+) m/z 631.9 $[M+H]^+$

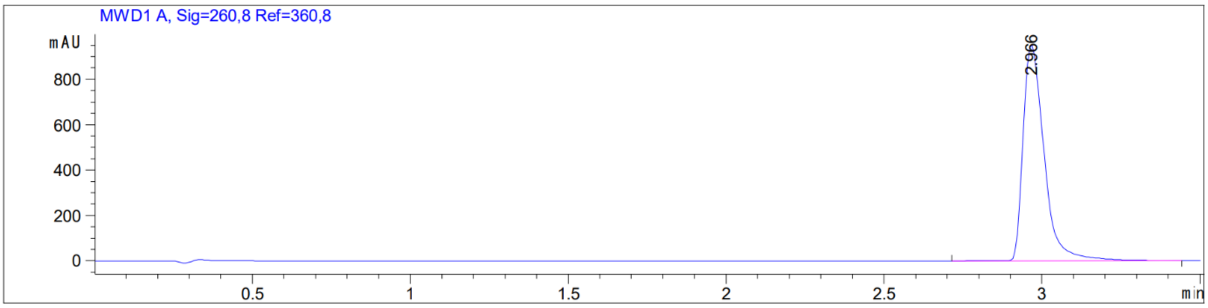

Synthesis of **$^{13}C_4$-OncoFAP-piperazine** (**7**)

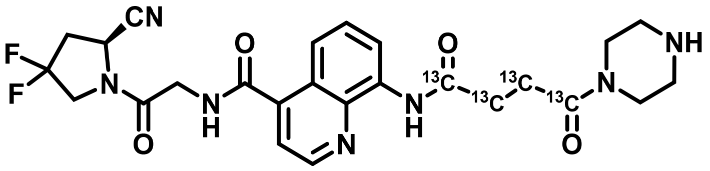

**$^{13}C_4$-OncoFAP-piperazine-Boc** (**6**) (20 mg, 0,03 mmol, 1 eq.) was dissolved in DCM (0.5 mL) and TFA (0.25 mL) was added dropwise. The mixture was stirred at room temperature for 3h then solvent was evaporated, and the crude was purified via RP flash chromatography (Büchi Sepacore equipped with FlashPure ID C18 irregular column) using a gradient of water + 0.1% FA/ACN + 0.1% FA 98:2 to 0:100 in 40 min. The desired fractions were collected and lyophilized to afford a colorless oil. (9 mg, 56%)

MS (ESI+) m/z 531.9 $[M+H]^+$

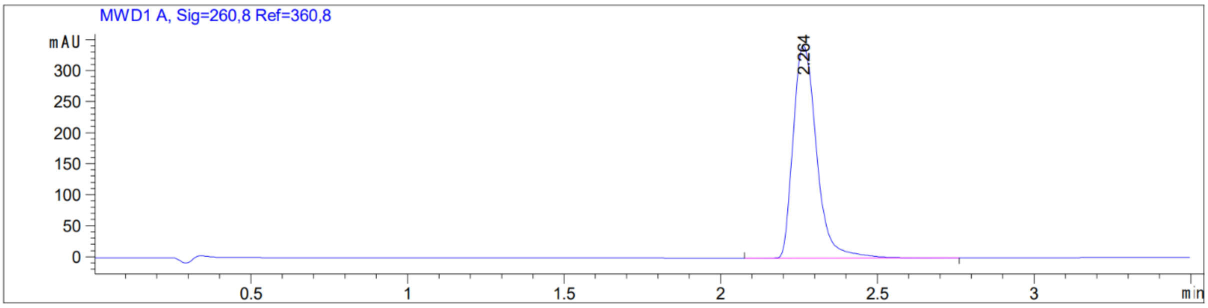

Synthesis of **$^{13}C_4$-OncoFAP-NOTA**

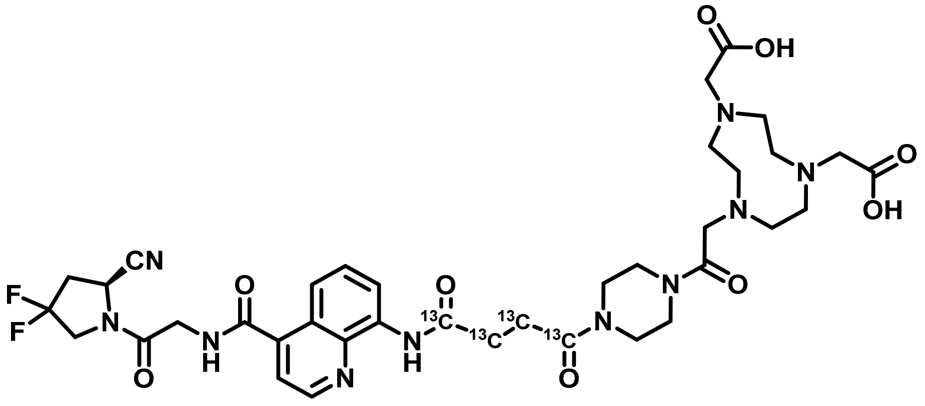

**$^{13}C_4$-OncoFAP-piperazine** (**7**) (2 mg, 0.005 mmol, 1 eq) and NOTA-NHS (4.5 mg, 0.01 mmol, 2 eq) were dissolved in DMF (50 µL). DIPEA (5 µL, 0.04 mmol, 4 eq) was added to the mixture and stirred for 1h at room temperature. The mixture was purified via RP-HPLC (Agilent 1200 series system equipped with Synergi MAX-RP C18 column using a gradient of 90:10 to 0:100 water + 0.1% TFA/ACN + 0.1% TFA in 12 min. The desired fractions were collected and lyophilized to afford a white solid. (2 mg, 25%)

MS (ESI+) m/z 816.7 $[M+H]^+$

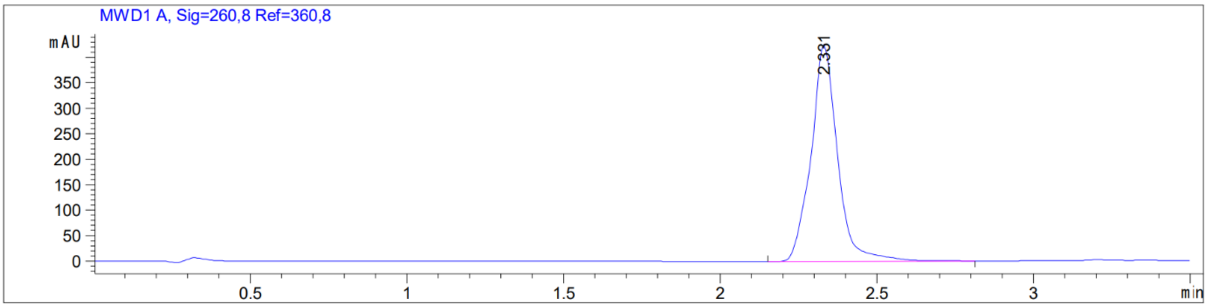

Synthesis of **[$^{nat}$F]AlF-$^{13}$C$_4$-OncoFAP-NOTA**

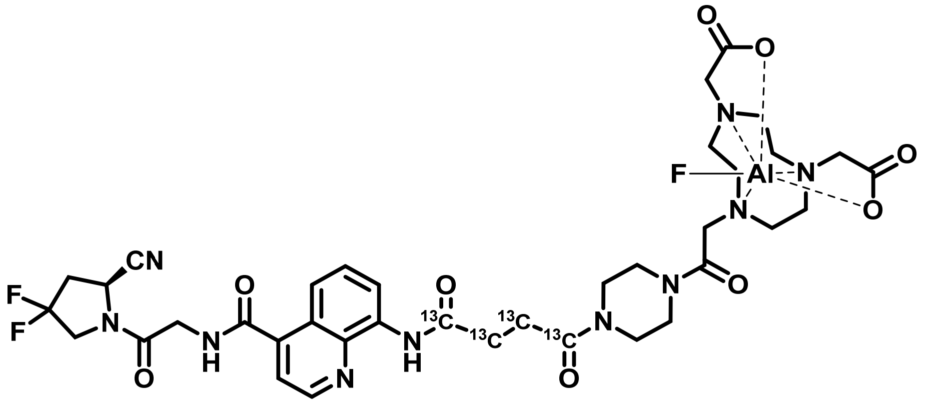

**$^{13}$C$_4$-OncoFAP-NOTA** (1 µmol) was dissolved in a mixture of DMSO (50 µL) and acetate buffer pH=4 (0.2 mL). $AlF_3$ (10 eq.) was added and the mixture was heated at 95°C for 15 min. Then the mixture was purified via RP-HPLC (Agilent 1200 series system equipped with Synergi MAX-RP C18 column) using a gradient of 95:5 to 0:100 water + 0.1% TFA/ACN + 0.1% TFA in 20 min. The desired fractions were collected and lyophilized to afford a white solid.

HRMS (ESI+) m/z 861.33401; theoretical m/z = 861.32671

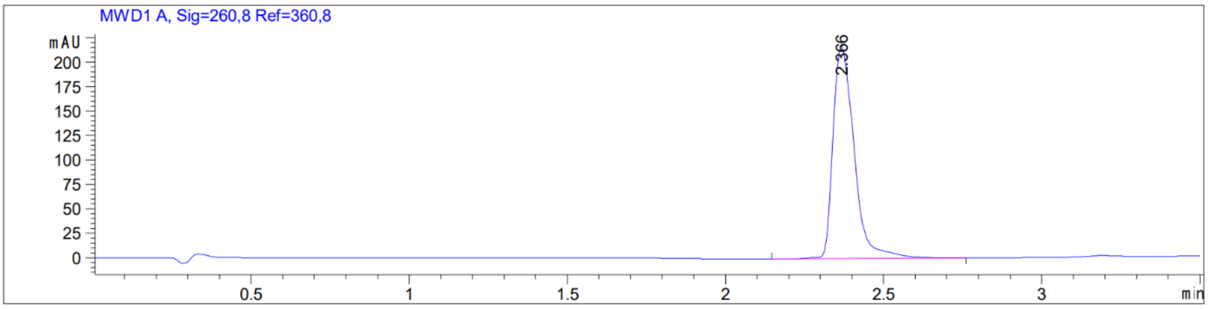

Synthesis of **OncoFAP-OSu** (**8**)

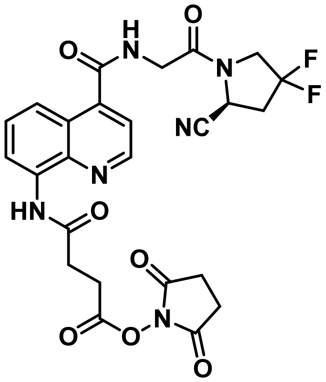

**OncoFAP-COOH** (**1**) (500 mg, 1.09 mmol, 1 eq), *N*-hydroxysuccinimide (188 mg, 1.63 mmol, 1.5 eq) and HATU (621 mg, 1.63 mmol, 1.5 eq) were dissolved in 5 mL of dry DMF, DIPEA (0.758 mL, 4.35 mmol, 4 eq.) was added dropwise. The reaction was stirred for 15 min, then diluted with DCM and washed with brine. The organic layer was dried over anhydrous Na2SO4, filtered and concentrated under reduced pressure. The dried crude was purified via column chromatography (DCM/MeOH 95:5 to 80:20) to obtain a pale yellow foam (600 mg, 99% yield).

MS (ESI+), m/z 557.2

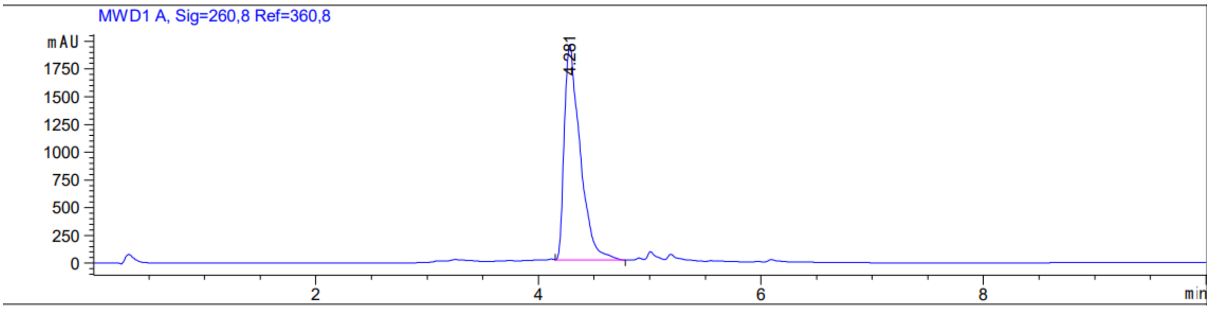

Synthesis of $^{13}C_6{}^{15}N_2$**BiOncoFAP-COOH** (**9**)

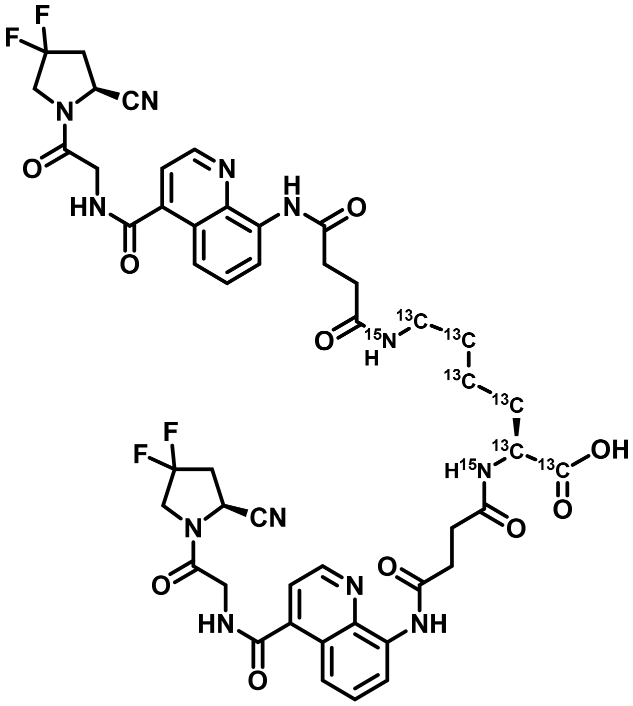

To a suspension of $^{13}C_6{}^{15}N_2$-L-Lysine (7 mg, 0.05 mmol, 1 eq.) in dry DMSO (0.7 mL), **OncoFAP-OSu** (**8**) (80 mg, 0.24 mmol, 3 eq.) was slowly added, followed by addition of DIPEA (0.08 mL, 0.48 mmol, 10 eq.). The mixture was stirred at room temperature for 30 minutes and then purified via RP-Flash Chromatography (Büchi Sepacore equipped with FlashPure ID C18 irregular column) using a gradient of water + 0.1% FA/ACN + 0.1% FA 98:2 to 0:100 in 40 min. The desired fractions were collected and lyophilized to afford an off-white solid. (18 mg, 37% yield).

MS (ESI+), m/z 1036.4

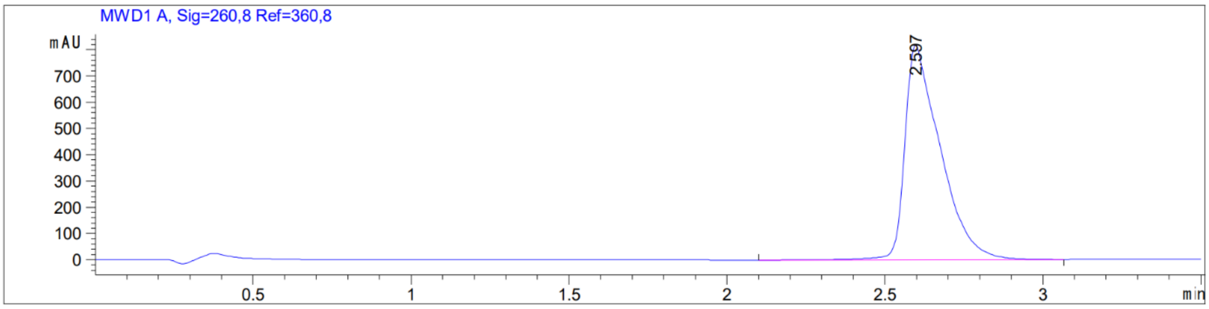

Synthesis of **$^{13}C_6{}^{15}N_2$BiOncoFAP-DOTAGA**

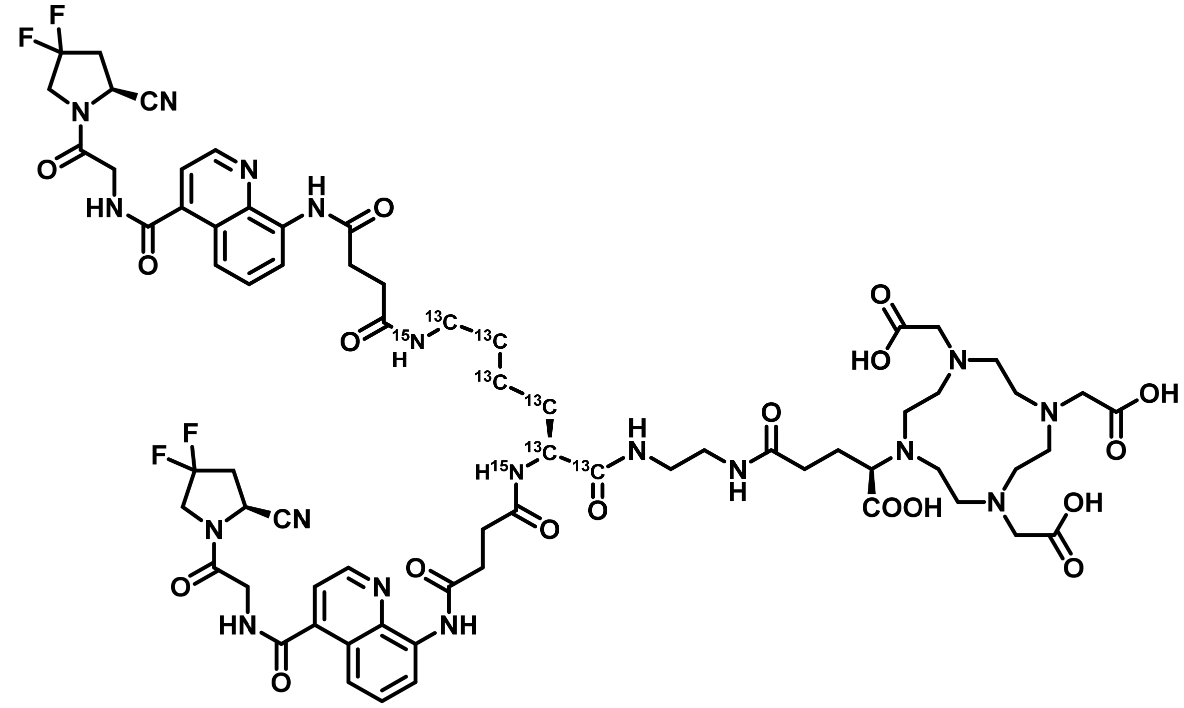

*N*-Hydroxysuccinimide (4 mg, 0.04 mmol, 2 eq.), HATU (8 mg, 0.02 mmol, 1.2 eq.) and DIPEA (0.02 mL, 0.09 mmol, 5 eq.) were added to a solution of **$^{13}C_6{}^{15}N_2$BiOncoFAP -COOH** (**9**) (18 mg, 0.02 mmol, 1 eq.) in dry DMF (0.5 mL). The reaction solution was stirred for 30 minutes at room temperature, then (*R*)-DOTAGA-$NH_2$ (18 mg, 0.04 mmol, 2 eq.) was added. The resulting mixture was stirred vigorously overnight, then diluted with milliQ water (0.5 mL) and purified via RP-HPLC (Agilent 1200 series system equipped with Synergi MAX-RP C18 column) using a gradient of 90:10 to 0:100 ACN + 0.1% TFA/water + 0.1% TFA in 12 min. The desired fractions were collected and lyophilized to afford a white solid. (14 mg, 52%)

MS (ESI+), m/z 1537.6

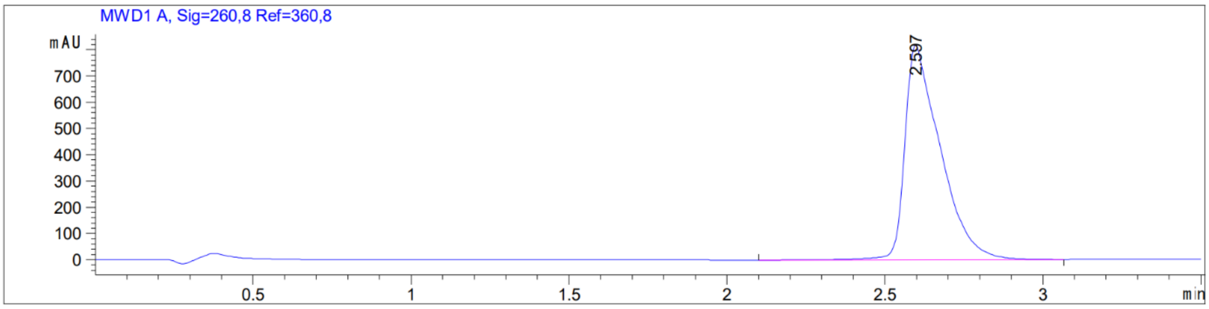

Synthesis of **[natLu]Lu-$^{13}C_6{}^{15}N_2$-BiOncoFAP-DOTAGA**

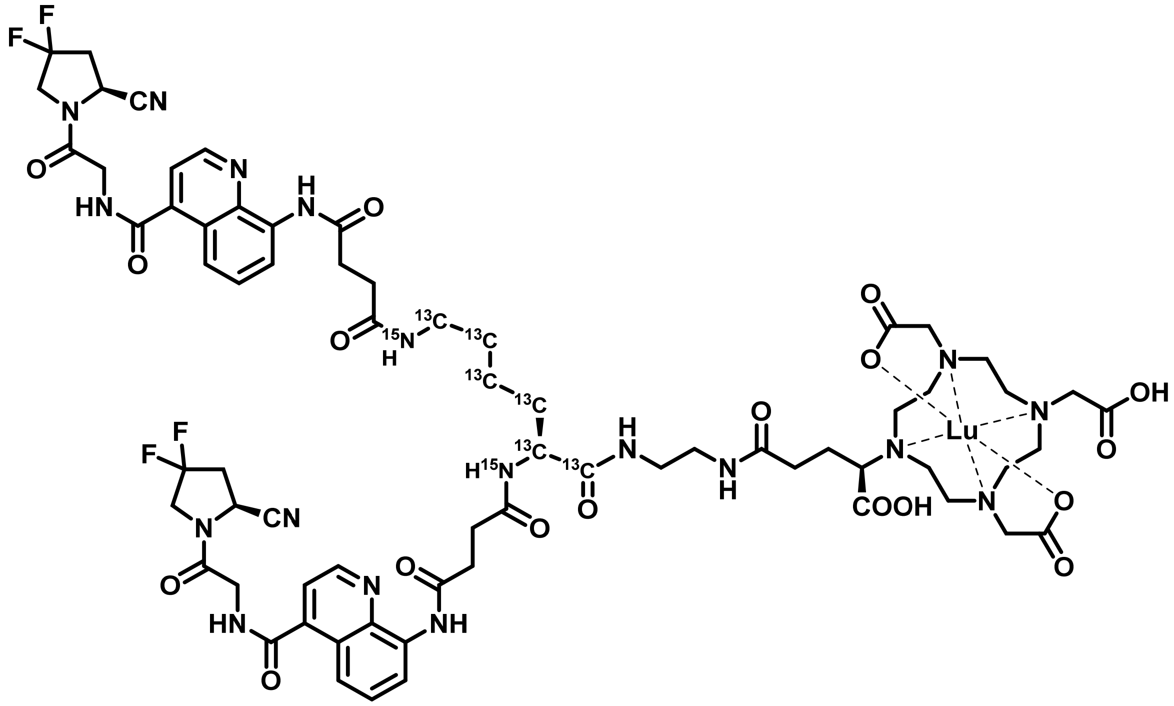

To a solution of **$^{13}C_6{}^{15}N_2$-BiOncoFAP-DOTAGA** (1 µmol) in 300 µL acetate buffer (aqueous solution, 1 M, pH 8), a freshly prepared solution of $LuCl_3$ hexahydrate (2 eq.) in 0.05N HCl (1.5 mL) was added. The resulting mixture was stirred at 95°C for 15 minutes, then purified via RP-HPLC (Agilent 1200 series system equipped with Synergi MAX-RP C18 column) using a gradient of 90:10 to 0:100 ACN + 0.1% TFA/water + 0.1% TFA in 12 min. The desired fractions were collected and lyophilized to afford a white solid.

HRMS (ESI+) m/z 1709.55655; theoretical m/z = 1709.55349

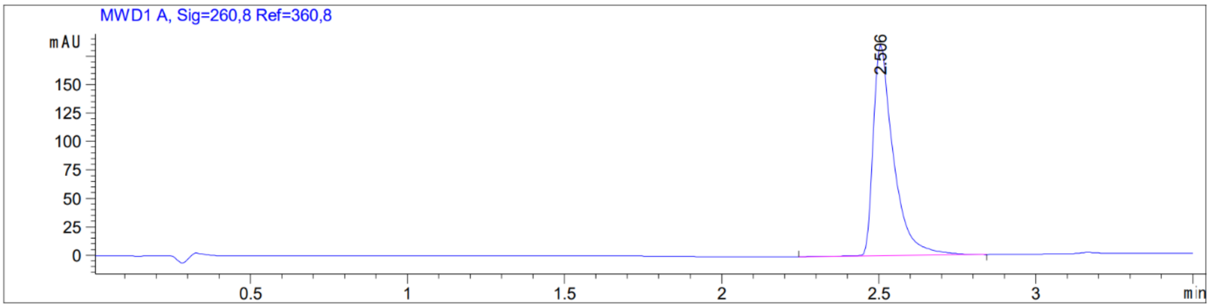

Synthesis of **[natGa]Ga-$^{13}C_6{}^{15}N_2$-BiOncoFAP-DOTAGA**

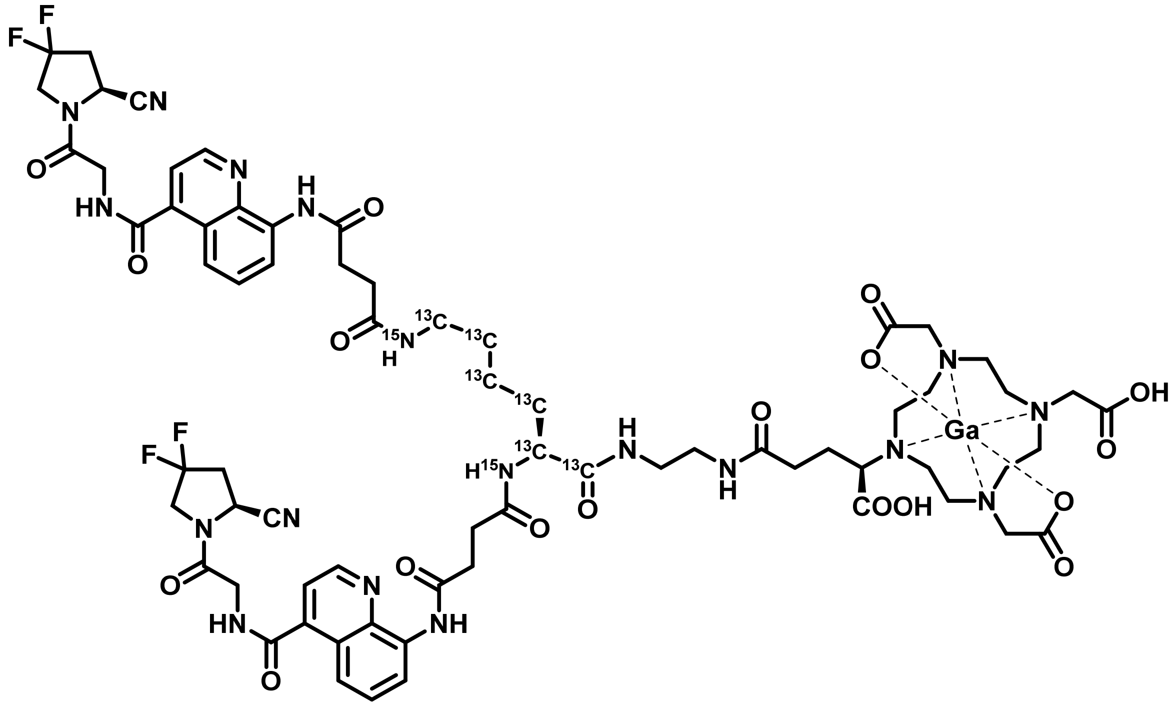

**$^{13}C_6{}^{15}N_2$-BiOncoFAP-DOTAGA** (1 µmol) was dissolved in acetate buffer, pH=4.5 (0.1 mL). Subsequently a solution of $GaCl_3$ (10 eq.) dissolved in 1N HCl (0.01 mL) was added. The reaction was stirred at 90°C for 10 min, then cooled down to room temperature. and purified via RP-HPLC (Agilent 1200 series system equipped with Synergi MAX-RP C18 column) using a gradient of 90:10 to 0:100 water + 0.1% TFA/ACN + 0.1% TFA in 16 min. The desired fractions were collected and lyophilized to afford a pale-yellow solid.

HRMS (ESI+) m/z 1603.53706 ($^{69}$Ga), 1605.53870 ($^{71}$Ga); theoretical m/z 1603.53828($^{69}$Ga), 1605.53741($^{71}$Ga)

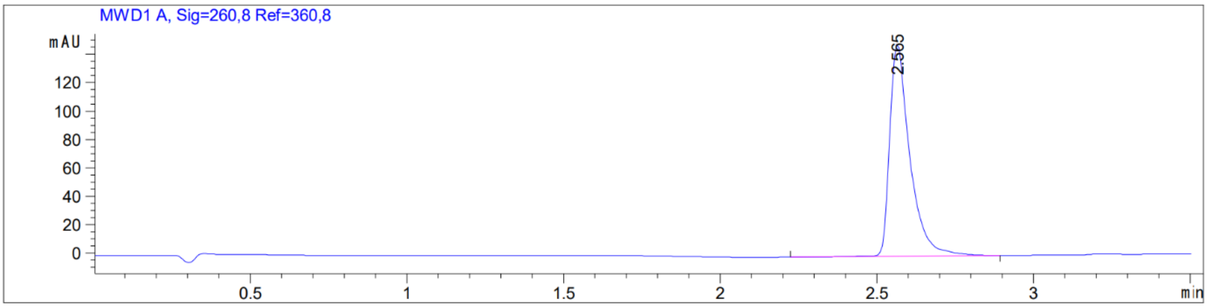

Synthesis of **[natGa]Ga-BiOncoFAP-DOTAGA**

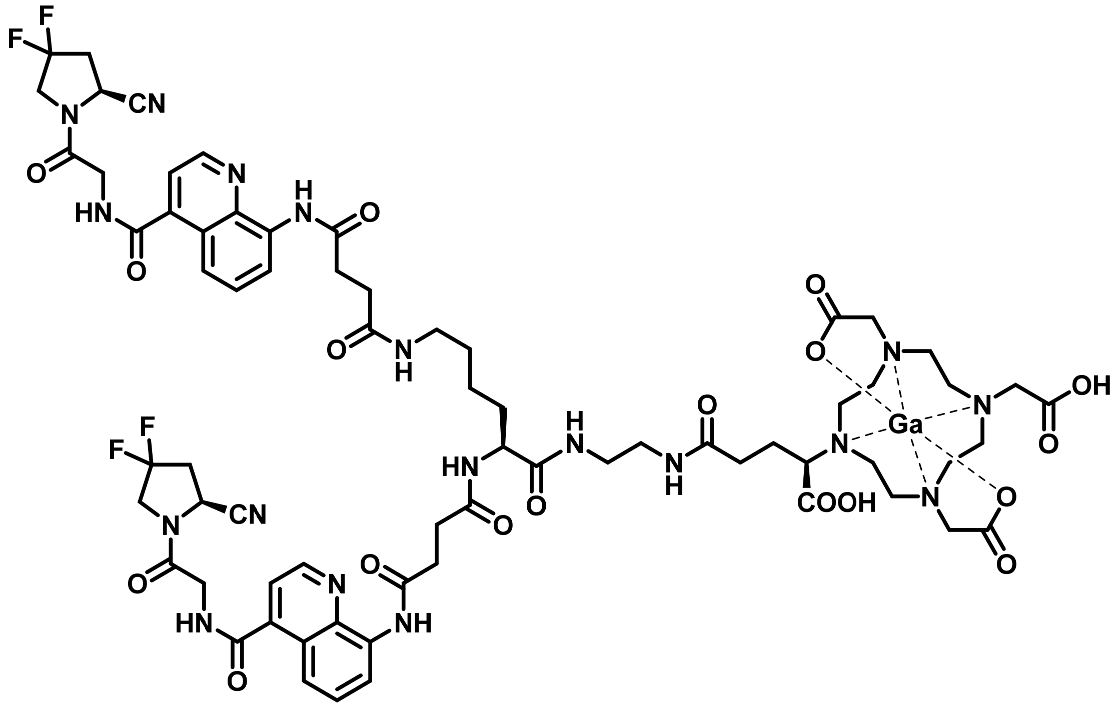

**BiOncoFAP-DOTAGA** (1 µmol) was dissolved in acetate buffer, pH=4.5 (0.1 mL). Subsequently a solution of $GaCl_3$ (10 eq.) dissolved in 1N HCl (0.01 mL) was added. The reaction was stirred at 90°C for 10 min, then cooled down to room temperature. and purified via RP-HPLC (Agilent 1200 series system equipped with Synergi MAX-RP C18 column) using a gradient of 90:10 to 0:100 water + 0.1% TFA/ACN + 0.1% TFA in 16 min. The desired fractions were collected and lyophilized to afford a pale-yellow solid.

HRMS (ESI+) m/z 1595.53054 ($^{69}$Ga), 1597.53223 ($^{71}$Ga) ; theoretical m/z 1595.52408($^{69}$Ga), 1597.52321($^{71}$Ga)

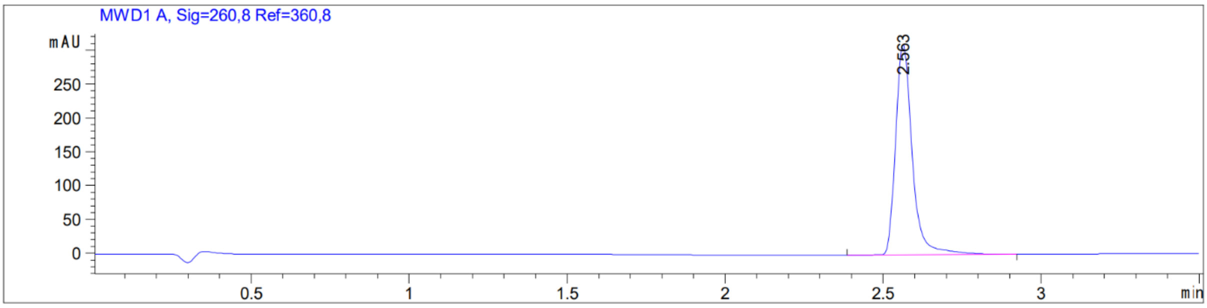

### Method Development

#### Quantitative analysis

To determine the single point calibration ratio, a fixed amount of analyte was added to a "blank" tumor tissue together with a fixed amount of the respective internal standard prior to tissue homogenization. Samples were then processed and analyzed with the nanoLC-HRMS following the same procedures described in the experimental section. Results are reported in table S1.

| Analyte | pmol injected in the MS | Ratio with IS |
|---|---|---|
| **[$^{nat}$Lu]Lu-OncoFAP-DOTAGA** | 1.5 | 1.01 ± 0.01 |
| **[$^{nat}$Lu]Lu-BiOncoFAP-DOTAGA** | 1.5 | 0.50 ± 0.01 |
| **[$^{nat}$Ga]Ga-OncoFAP-DOTAGA** | 1.5 | 2.06 ± 0.05 |
| **[$^{nat}$Ga]Ga-BiOncoFAP-DOTAGA** | 1.5 | 1.47 ± 0.01 |
| **[$^{nat}$F]AlF-OncoFAP-NODAGA** | 0.39 | 3.7 ± 0.08 |
| **Al-OncoFAP-NODAGA** | 1.62 | 0.21 ± 0.01 |
| **AlOH-OncoFAP-NODAGA** | 0.96 | 50.75 ± 0.55 |
| **[$^{nat}$F]AlF-OncoFAP-NOTA** | 3.0 | 0.27 ± 0.01 |

**Table S1**: Ratios between MS areas of fixed amounts of analytes and corresponding internal standards were calculated and used as external calibration points for quantification of *in vivo* biodistribution experiments. Values are expressed as mean ± sd, n = 4.

**Matrix Effect and Recovery in tumor**

“Blank” tumors were processed following the same procedures described in the experimental section. To evaluate the matrix effect, 30 μL of a solution containing all analytes at a final concentration of 1 μM was used to resuspend “blank” tumor tissues. Analytes in matrix and analytes in aqueous solution were then analyzed by LC-HRMS using the same methods reported in the experimental section. Matrix effect was calculated with equation S1**.**

**Equation S1** $$\text{Matrix Effect} = 1 - \frac{\text{AUC}_{\text{after sample prep}}}{\text{AUC}_{\text{water}}} \times 100$$

To evaluate the recovery, 50 μL of a solution containing all the analytes at a final concentration of 1.2 μM was added during the sample preparation of untreated “blank” tumors specimens before the homogenization. Tumors were then processed as reported in the experimental section. Samples were analyzed by nanoLC-HRMS as previously described. Recovery was calculated with equation S2.

**Equation S2** $$\text{Recovery} = \frac{\text{AUC}_{\text{before sample prep}}}{\text{AUC}_{\text{after sample prep}}} \times 100$$

| Analyte | Matrix Effect | Recovery |
|---|---|---|
| **[$^{nat}$Lu]Lu-OncoFAP-DOTAGA** | 99.14 ± 0.02 | 49.11 ± 7.14 |
| **[$^{nat}$Lu]Lu-BiOncoFAP-DOTAGA** | 92.06 ± 1.70 | 48.75 ± 19.89 |
| **[$^{nat}$Ga]Ga-OncoFAP-DOTAGA** | 98.80 ± 0.52 | 53.07 ± 29.46 |
| **[$^{nat}$Ga]Ga-BiOncoFAP-DOTAGA** | 92.84 ± 1.30 | 38.96 ± 4.42 |
| **[$^{nat}$F]AlF-OncoFAP-NODAGA** | 64.08 ± 2.47 | 44.51 ± 8.80 |
| **Al-OncoFAP-NODAGA** | 59.05 ± 5.66 | 45.45 ± 8.46 |
| **AlOH-OncoFAP-NODAGA** | 46.91 ± 8.77 | 50.26 ± 12.31 |
| **[$^{nat}$F]AlF-OncoFAP-NOTA** | 58.98 ± 4.55 | 50.49 ± 3.48 |

**Table S2**: Matrix effect and recovery calculated for each single analyte used in the study. Values are expressed as mean ± sd, n = 3.

### Tables

| %ID/g | [natLu]Lu-OncoFAP-DOTAGA | | | [natLu]Lu-BiOncoFAP-DOTAGA | | |
|---|---|---|---|---|---|---|
| | Mouse 1 | Mouse 2 | Mouse 3 | Mouse 1 | Mouse 2 | Mouse 3 |
| **Tumor** | 21.36 | 23.46 | 21.19 | 27.39 | 29.69 | 23.35 |
| **Plasma** | 0.40 | 0.55 | 0.42 | 2.11 | 2.80 | 1.56 |
| **Spleen** | 0.12 | 0.10 | 0.10 | 0.56 | 0.67 | 0.47 |
| **Kidney** | 0.57 | 1.07 | 0.96 | 3.32 | 4.96 | 2.63 |
| **Heart** | 0.03 | 0.15 | 0.04 | 0.39 | 0.88 | 0.34 |
| **Lung** | 0.08 | 0.35 | 0.16 | 1.60 | 2.16 | 1.32 |
| **Liver** | 2.57 | 3.06 | 2.91 | 1.46 | 2.07 | 1.18 |

**Table S3**: Calculated %ID/g of [natLu]Lu-OncoFAP-DOTAGA, and [natLu]Lu-BiOncoFAP-DOTAGA in biodistribution experiments performed on mice bearing HT-1080.hFAP tumors sacrificed 1h after intravenous administration (250 nmol/kg).

| %ID/g | [natLu]Lu-OncoFAP-DOTAGA vs [177Lu]Lu-OncoFAP-DOTAGA | | [natLu]Lu-BiOncoFAP-DOTAGA vs [177Lu]Lu-BiOncoFAP-DOTAGA | | [natGa]Ga-OncoFAP-DOTAGA vs [natLu]Lu-OncoFAP-DOTAGA | | [natGa]Ga-BiOncoFAP-DOTAGA vs [natLu]Lu-BiOncoFAP-DOTAGA | |
|---|---|---|---|---|---|---|---|---|
| | **p value** | **<0.05** | **p value** | **<0.05** | **p value** | **<0.05** | **p value** | **<0.05** |
| **Tumor** | 0.300 | False | 0.578 | False | 0.039 | False | 0.495 | False |
| **Plasma** | 0.261 | False | 0.031 | False | 0.082 | False | 0.866 | False |
| **Spleen** | 0.001 | **True** | 0.413 | False | 0.916 | False | 0.051 | False |
| **Kidney** | 0.006 | False | 0.598 | False | 0.416 | False | 0.240 | False |
| **Heart** | 0.167 | False | 0.897 | False | 0.422 | False | 0.666 | False |
| **Lung** | 0.019 | False | 0.649 | False | 0.711 | False | 0.272 | False |
| **Liver** | 0.031 | False | 0.927 | False | 0.001 | **True** | 0.042 | **True** |

**Table S4**: p values of multiple t-test are shown comparing %ID/g in each organ for i) [natLu]Lu-OncoFAP-DOTAGA and [177Lu]Lu-OncoFAP-DOTAGA; ii) [natLu]Lu-BiOncoFAP-DOTAGA and [177Lu]Lu-BiOncoFAP-DOTAGA; iii) [natGa]Ga-OncoFAP-DOTAGA and [natLu]Lu-OncoFAP-DOTAGA; iiii) [natGa]Ga-BiOncoFAP-DOTAGA and [natLu]Lu-BiOncoFAP-DOTAGA. %ID/g of [177Lu]Lu-OncoFAP-DOTAGA and %ID/g of [177Lu]Lu-BiOncoFAP-DOTAGA were measured by radioactivity using a gamma-counter[3] while all other compounds were measured by MS.

| %ID/g | [natGa]Ga-OncoFAP-DOTAGA | [natGa]Ga-BiOncoFAP-DOTAGA |
|---|---|---|

| | Mouse 1 | Mouse 2 | Mouse 3 | Mouse 1 | Mouse 2 | Mouse 3 |
|---|---|---|---|---|---|---|
| **Tumor** | 13.83 | 19.01 | 11.02 | 36.00 | 22.72 | 31.43 |
| **Plasma** | 0.33 | 0.36 | 0.34 | 1.31 | 1.41 | 3.35 |
| **Spleen** | 0.03 | 0.05 | 0.22 | 0.25 | 0.30 | 0.45 |
| **Kidney** | 0.74 | 0.67 | 0.77 | 1.79 | 1.43 | 3.65 |
| **Heart** | 0.10 | 0.11 | 0.11 | 0.23 | 0.19 | 0.81 |
| **Lung** | 0.17 | 0.02 | 0.27 | 0.79 | 0.80 | 1.85 |
| **Liver** | N.D. | 0.20 | 0.25 | 0.46 | 0.47 | 0.98 |

**Table S5**: Calculated %ID/g of [natGa]Ga-OncoFAP-DOTAGA, and [natGa]Ga-BiOncoFAP-DOTAGA in biodistribution experiments performed on mice bearing HT-1080.hFAP tumors sacrificed 1h after intravenous administration (250 nmol/kg).

| **%ID/g** | **Al-OncoFAP-NODAGA** | | | **AlOH-OncoFAP-NODAGA** | | | **[natF]AlF-OncoFAP-NODAGA** | | |
|---|---|---|---|---|---|---|---|---|---|
| | Mouse1 | Mouse2 | Mouse3 | Mouse1 | Mouse2 | Mouse3 | Mouse1 | Mouse2 | Mouse3 |
| **Tumor** | 2.73 | 5.96 | 5.33 | 0.02 | 0.02 | 0.03 | 0.29 | 0.65 | 0.53 |
| **Plasma** | 0.1 | 0.19 | 0.09 | 0.00 | 0.01 | 0.00 | 0.01 | 0.01 | 0.01 |
| **Spleen** | 0.01 | 0.01 | 0.01 | 0.00 | 0.00 | 0.01 | 0.00 | 0.00 | 0.01 |
| **Kidney** | 0.21 | 0.12 | 0.16 | 0.11 | 0.07 | 0.15 | 0.05 | 0.03 | 0.04 |
| **Heart** | 0.05 | 0.01 | 0.02 | 0.01 | 0.00 | 0.00 | 0.01 | 0.00 | 0.00 |
| **Lung** | 0.08 | 0.05 | 0.08 | 0.02 | 0.02 | 0.02 | 0.01 | 0.01 | 0.00 |
| **Liver** | 0.58 | 0.62 | 0.5 | 0.01 | 0.12 | 0.03 | 0.02 | 0.00 | 0.03 |

**Table S6**: Calculated %ID/g of of the three different molecular species, Al-OncoFAP-NODAGA, AlOH-OncoFAP-NODAGA and [natF]AlF-OncoFAP-NODAGA, in biodistribution experiments performed on mice bearing SK-RC-52.hFAP tumors tumors sacrificed 2h after intravenous administration (500 nmol/kg).

| **%ID/g** | **OncoFAP-NODAGA** | **[natF]AlF-OncoFAP-NOTA** |
|---|---|---|

| | Mouse 1 | Mouse 2 | Mouse 3 | Mouse 1 | Mouse 2 | Mouse 3 |
|---|---|---|---|---|---|---|
| **Tumor** | 3.04 | 6.64 | 5.90 | 13.35 | 13.54 | 11.62 |
| **Plasma** | 0.11 | 0.21 | 0.10 | 0.37 | 0.30 | 0.32 |
| **Spleen** | 0.01 | 0.02 | 0.03 | 0.08 | 0.04 | 0.01 |
| **Kidney** | 0.37 | 0.21 | 0.35 | 0.55 | 0.50 | 0.39 |
| **Heart** | 0.06 | 0.01 | 0.03 | 0.10 | 0.10 | 0.04 |
| **Lung** | 0.11 | 0.07 | 0.10 | 0.29 | 0.21 | 0.17 |
| **Liver** | 0.61 | 0.74 | 0.55 | 0.16 | 0.16 | 0.16 |

**Table S7**: Calculated %ID/g of OncoFAP-NODAGA (sum of three different molecular species), and [$^{nat}$F]Al-F-OncoFAP-NOTA in biodistribution experiments performed on mice bearing SK-RC-52.hFAP tumors tumors sacrificed 2h after intravenous administration (500 nmol/kg).

### Figures

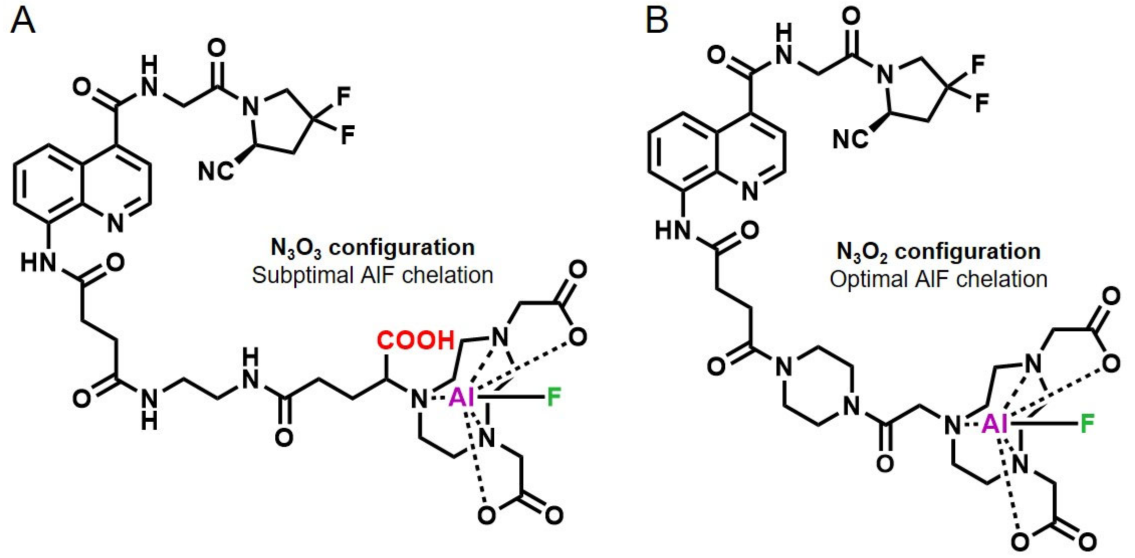

**Figure S1**: A) Chemical structure of [natF]AlF-OncoFAP-NODAGA, showing the $N_3O_3$ configuration which is unfavorable to AlF chelation; B) chemical structure of [natF]AlF-OncoFAP-NOTA, showing the $N_3O_2$ conformation, favorable to AlF chelation.

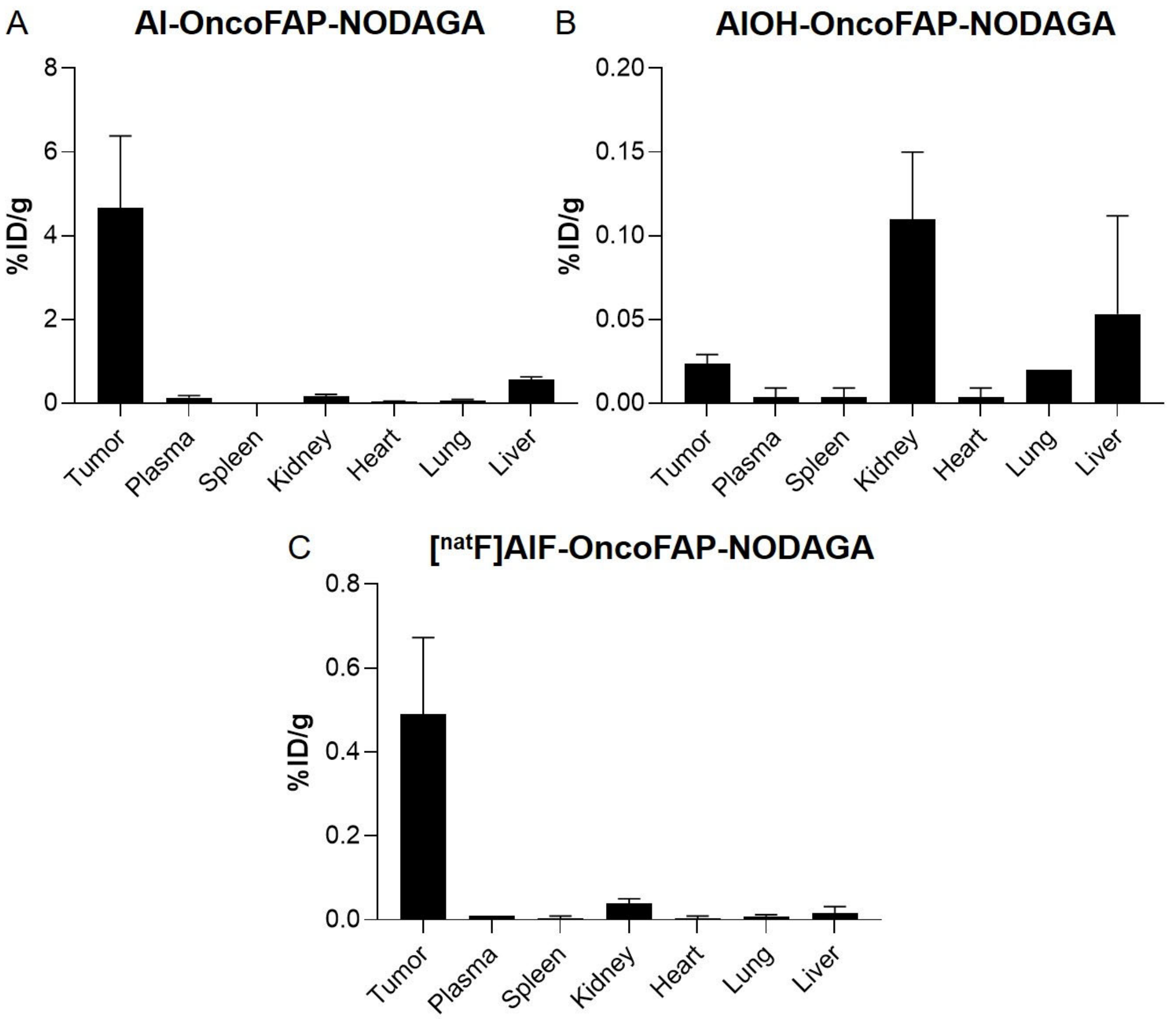

**Figure S2**: MS biodistribution results of A) Al-OncoFAP-NODAGA; B) AlOH-OncoFAP-NODAGA; C) [natF]AlF-OncoFAP-NODAGA. Mice bearing SK-RC-52.hFAP tumors were sacrificed 2 h after intravenous administration at a dose of 500 nmol/kg.
